# Global Analysis of Aggregation Determinants in Small Protein Domains

**DOI:** 10.1101/2025.11.11.687847

**Authors:** Cydney M. Martell, Kamil K. Gebis, Han My Van, Yulia M. Gutierrez, Michelle D. Jung, Jeffrey N. Savas, Gabriel J. Rocklin

## Abstract

Protein aggregation is an obstacle for engineering effective recombinant proteins for biotechnology and therapeutic applications. Predicting protein aggregation propensity remains challenging due to the complex interplay of sequence, structure, environmental factors, and external stress conditions, particularly for globular proteins. To understand the determinants of aggregation and improve its prediction, we quantified insoluble aggregation following high temperature and acidic stress in custom libraries of small protein domains (40-72 amino acids) using a high-throughput, in vitro, mass spectrometry-based method. In total, we quantified aggregation for 18,987 small protein domains, revealing diverse stress-dependent aggregation phenotypes that were consistent in different library contexts. We also found that aggregation measurements on individually purified proteins strongly correlated with high-throughput mixed-pool data (Pearson’s r = 0.65-0.79), supporting the use of multiplexed approaches to study aggregation. Using machine learning, we identified sequence and structural features that correlate with aggregation and fine-tuned the protein language model SaProt, which explained 43-55% of the observed variation in a held-out test set of unrelated protein domains. Our model shows promising utility for engineering aggregation-resistant proteins, and our dataset serves as an important resource for developing improved models of protein aggregation.

## Main

Protein aggregation is a major challenge for the use of proteins in therapeutic and biotechnology applications. Despite its importance, efforts to engineer aggregation-resistant proteins remain largely empirical, due to the lack of predictive models that generalize across diverse sequences and conditions. Aggregation is driven by a combination of intrinsic sequence features (Chiti et al., 2003; Navarro & Ventura, 2022) and extrinsic factors such as pH, temperature, concentration, and agitation (Roberts, 2014; Schvartz et al., 2023; W. Wang et al., 2010). While many computational tools exist to predict aggregation prone regions, most are based on amyloid-β peptide aggregation behavior (Conchillo-Solé et al., 2007; Keresztes et al., 2021; Maurer-Stroh et al., 2010; Prabakaran et al., 2021), physicochemical properties (Fernandez-Escamilla et al., 2004; Orlando et al., 2020; Tartaglia & Vendruscolo, 2008; Zibaee et al., 2007), or β-sheet pairing energies derived from protein structures (Trovato et al., 2007). As a result, they often fail to predict key residues influencing aggregation in globular proteins (Guthertz et al., 2022; Meric et al., 2021). This highlights the need for improved models based on experimental measures across diverse globular proteins and conditions.

Existing experimental datasets have not yet led to robust predictive models of globular protein aggregation. Many studies have focused on deep investigations of individual proteins of interest (Ebo et al., 2020; Guthertz et al., 2022), limiting their generalizability. Large-scale cell-free studies have measured soluble expression of the *E. coli* proteome (Niwa et al., 2009) and hundreds of eukaryotic proteins (Uemura et al., 2018). Other high-throughput studies have measured in vivo aggregation of the human proteome in response to heat shock (Määttä et al., 2020) and thermal stability for tens of thousands of proteins in cells (Jarzab et al., 2020). However, these in vivo results are complicated by the complexity of the cellular environment, wide variation in the abundances of different cellular proteins, and heterogeneity in proteoforms (e.g., from post-translational modifications) that make it difficult to link chemical structure to aggregation phenotypes. Some large-scale studies focus specifically on amyloid aggregation (Arutyunyan et al., 2025; Seuma et al., 2022; M. Thompson et al., 2025), but this represents only one of the several mechanisms by which proteins can aggregate. Overall, existing datasets do not provide quantitative, high-throughput measurements of aggregation behavior across a wide range of sequences under controlled experimental conditions.

To address these limitations, we measured aggregation of small protein domains in custom libraries under controlled stress conditions using a quantitative in vitro mass spectrometry (MS) assay. We synthesized large mixtures of small protein domains from custom DNA oligo pools (Ferrari et al., 2025) and measured aggregation propensity within these complex mixtures by quantifying how much of each protein remains soluble after stress treatment. While mass spectrometry has previously been applied to study protein solubility and oligomeric state (Feldman et al., 2025; Jarzab et al., 2020; Määttä et al., 2020), our method of measuring stress-induced aggregation introduces several innovations. First, by focusing on small protein domains (<10 kDa), we leverage simplified model systems that are ideal for directly connecting sequence and structural features to aggregation behavior. Second, constructing libraries using synthetic DNA provides full control over sequence content, allowing inclusion of both natural and *de novo* designed domains and enabling systematic analysis of small, structurally defined domains, in contrast to complex cellular proteomes. These domains could also be re-designed and encoded in oligo pools for iterative testing of our models at scale. Third, our in vitro platform avoids complications from cellular factors such as abundance, localization, chaperone activity, and proteoform heterogeneity (e.g., post translational modifications). Notably, we also measured the global folding stability of these domains using the cDNA display proteolysis method developed in our lab (Tsuboyama et al., 2023), enabling us to investigate how folding stability impacts aggregation.

By quantifying the depletion of soluble small protein domains following exposure to thermal or acidic stress, we measured insoluble aggregation for 18,987 small protein domains (40-72 amino acids). These data revealed a range of stress-dependent aggregation phenotypes across domains. Notably, we found that aggregation measurements for individually purified proteins correlated with measurements obtained from the high-throughput mixed-pool experiments. Using these data, we identified sequence and structural features that govern stress-induced aggregation and trained predictive models that generalize to proteins outside the training set. Our model shows promise for engineering aggregation resistance in proteins and our dataset serves as an important resource for future machine learning models.

## Results

### High-throughput approach to quantify stress-induced aggregation

To measure aggregation across a diverse range of proteins, we assayed two types of libraries: the Metagenomic libraries and the Protein Family library. The Metagenomic libraries consist of structurally diverse small domains chosen from the MGnify microbiome sequence database (Richardson et al., 2023). These domains were randomly subdivided into two sub-libraries (Metagenomic 1 and Metagenomic 2) to test the impact of library context on aggregation measurements, with a subset of proteins included in both libraries. The Protein Family library consists of natural domains from the Pfam database (Finn et al., 2014) and *de novo* designed folds (Kim et al., 2022; Rocklin et al., 2017) (Fig. 1A). Despite their small size, these domains fold into diverse tertiary structures (Fig. 1A). We constructed these libraries to achieve a high degree of diversity in biophysical features (e.g., hydrophobicity and pI) in each library and to ensure that each domain had several unique predicted tryptic peptides suitable for bottom-up MS detection, avoiding the need for peptide barcodes (Fig. 1B). Additionally, we selected proteins with high sequence diversity to promote generalizability of our conclusions, with the average sequence identity between nearest-neighbor proteins ranging from 38–50% in each library (Fig. 1B).

**Figure 1.**
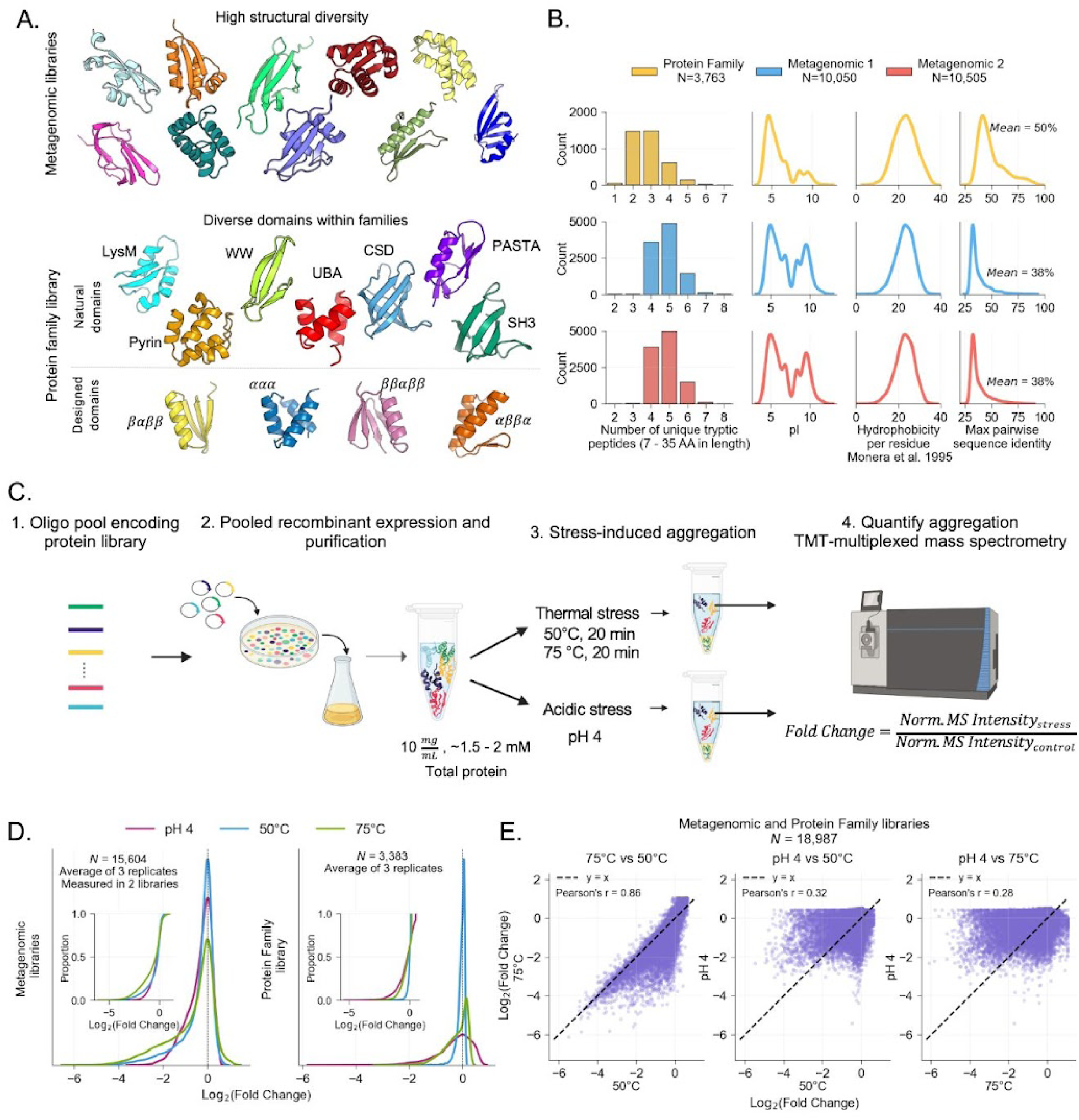
High-throughput in vitro approach to measure stress-induced aggregation of small protein domains. (A) Representative Alphafold 2-predicted structures of the domains selected in Metagenomic and Protein Family libraries. (B) Distributions of protein features across domains in each library. Proteins were selected to have unique tryptic peptides for bottom-up mass spectrometry and a diverse range of features. Hydrophobicity was calculated using the scale from Monera et al. (1995), based on reversed-phase HPLC retention times of amphipathic α-helical peptides with single amino acid substitutions. pI was calculated using pK values from Bjellqvist et al. (1993 & 1994). (C) Mass spectrometry-based quantification of protein aggregation. Selected domains are encoded in oligo pools, expressed and purified as a pooled mixture and concentrated to 10 mg/ml. Aggregation is induced by high temperature stress (50 °C, 75 °C) or acidic stress (pH 4), after which aggregates are removed by centrifugation. Remaining soluble protein is quantified by TMT-multiplexed mass spectrometry and compared to a control sample (room temperature for thermal stress and pH 7 for acidic stress). (D) Distributions of log_2_(Fold Change) for each stress condition (50 °C: blue, 75 °C: green, pH 4: pink) for the Metagenomic Libraries (left) and Protein Family library (right). Data represents an average of 3 replicates. The dashed line represents x = 0, negative values indicate aggregation-prone proteins. The inset shows the same data represented as an empirical cumulative distribution function plot. (E) Correlation of log_2_(Fold Change) between stress conditions for all quantified protein domains. The dashed line indicates y=x.

We ordered DNA oligo pools encoding the selected domains in each library (∼3,000 to 10,000 domains per library) and used pooled recombinant expression to synthesize the mixed protein libraries, effectively creating a custom proteome (Fig. 1C; Ferrari et al., 2025). Prior to aggregation, we isolated the monomeric fraction of each library using size-exclusion chromatography and concentrated the sample to 10 mg/ml (Fig. 1C, Fig. S1A-B). We then induced aggregation with thermal (50 °C or 75 °C for 20 min) or acidic (pH 4 for 10 min) stress and removed insoluble aggregates by centrifugation. Alongside each stress condition, we included control samples (room temperature or pH 7) to account for differences in starting abundance and MS ionization efficiency between domains. We analyzed the soluble fraction from stressed and control samples using Tandem Mass Tag (TMT)-multiplexed bottom-up mass spectrometry (Li et al., 2020; A. Thompson et al., 2003). Aggregation propensity was calculated for each protein as the ratio of normalized MS intensity in stressed versus control samples (Fig. 1C). We represent aggregation on a log_2_ scale, where more negative values indicate greater protein aggregation (decrease in soluble protein abundance). For example, an aggregation propensity of −4 means the protein’s relative abundance after stress is 1/16 of its abundance in the control sample. Positive values do not indicate a true increase in relative abundance but reflect MS normalization effects (see methods). Using this approach, we identified 19,063 domains across all libraries and successfully measured thermal- and acid-induced aggregation for 18,987 unique domains (∼82% of the domains encoded in the DNA libraries; Fig. 1D, Fig. S2A).

Protein aggregation is known to depend strongly on concentration, with higher concentrations often promoting aggregation (Ciryam et al., 2015; Siddiqui & Naeem, 2018). In our pooled experiments, however, we can’t control the concentration of every individual protein in the mixture, as they likely express at different levels. At the library level, we observed reduced aggregation at lower protein concentrations compared to higher concentrations, although this effect decreased above approximately 7.5-10 mg/ml (Fig. S1D). We therefore assayed all libraries at a total protein concentration of 10 mg/ml, which produced similar total protein loss to higher concentrations (Fig. S1D). Aggregation measurements should be interpreted within the context of this concentration, as the behavior of individual proteins may differ at other concentrations.

Aggregation measurements were reproducible. Each stress experiment was performed in triplicate starting from the same protein preparation. Normalized TMT intensity correlated very well across the three replicates in each stress (Pearson’s r > 0.9 for most conditions; Fig. S6A) and aggregation measurements showed a low standard deviation for the majority of proteins (median standard deviation of log_2_(fold change) = 0.19-0.27; Fig. S6B), showing the robustness of our assay.

Some domains were not detected by MS, even in the unstressed sample, likely due to insoluble expression or peptide properties that reduce detectability. Domains not identified by MS tend to be more hydrophobic and have lower experimental folding stability, as measured by cDNA display proteolysis, indicating some domains may not express solubly (Fig. S2B). Additionally, domains that were not identified also tend to have fewer predicted unique tryptic peptides, likely reflecting poor MS detectability (Fig. S2C). Consequently, our experimental libraries are slightly biased toward solubly expressing domains and ones that have optimal properties for MS detection. Despite these biases, the domains in our assay still represent 82% of the ordered domains across all 3 libraries and span a large range across protein features (Fig. S2D).

### Domains show a broad range of aggregation phenotypes

We observed a range of aggregation phenotypes, spanning from domains that were highly aggregation-prone (relative abundance dropped by >80%) to those that remained largely soluble after exposure to stress. (Fig. 1D). Overall, domains in the Protein Family library aggregated less under high temperature stress than the Metagenomic domains, while both exhibited similar levels of aggregation at low pH, consistent with total protein quantification (Fig. 1D, Fig. S1C). Aggregation propensities were strongly correlated between 50 °C and 75 °C (Pearson’s r = 0.86), though some proteins aggregated only after exposure to the higher temperature (aggregation never substantially decreased at 75°C compared to 50 °C). In contrast, aggregation from thermal and acidic conditions were weakly correlated (Pearson’s r = 0.28), with many domains aggregating in only one condition (Fig. 1E). Overall, sequence similarity trended with similarity in aggregation phenotypes under each stress condition (Fig. 2A), indicating that our measurements capture meaningful sequence-phenotype relationships in protein aggregation.

**Figure 2.**
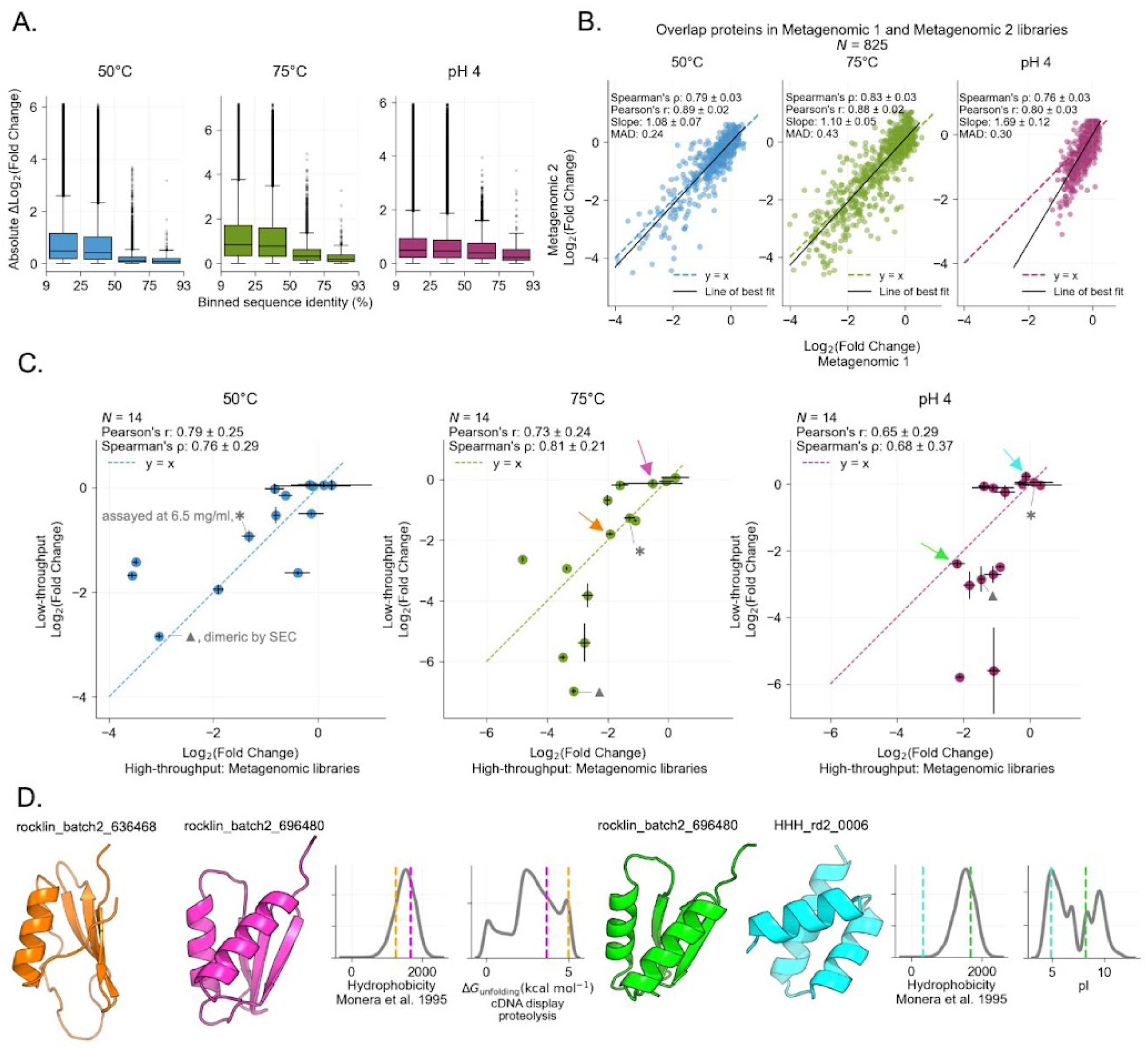
Aggregation measurements in pooled libraries are consistent in several library contexts and correlate well with measurements on individual proteins. (A) Relationship between sequence identity and aggregation phenotype for each stress (50 °C (blue, left), 75 °C (green, middle), pH 4 (pink, right)). A higher Δlog_2_(fold change) represents a larger difference in aggregation phenotype between protein pairs. Boxplots show the distribution for every protein pair in each of the three libraries binned by pairwise sequence identity (Number of pairs in each bin: 9-25%, N= 8.5×10^7^; 25-50%, N = 1.5×10^7^; 50-75%, N= 4.9×10^3^; 75-100%, N= 352). Boxplots show the median (center line), 25th and 75th percentiles (box edges), and whiskers extending to 1.5x the interquartile range. (B) Correlation of aggregation propensity measured for 825 proteins in two different libraries: Metagenomic 1 and Metagenomic 2. Metrics are represented as values ± half 95% CI from bootstrapping. (C) Comparison of high-throughput measurements and aggregation measured on individual proteins. For the low-throughput assay, proteins were individually purified, concentrated to 10 mg/ml, and subjected to the same stress-induced aggregation protocol used in the high-throughput approach. Aggregation was quantified by measuring the loss of soluble protein using a BCA assay on the supernatant, compared to the unstressed control. Error bars represent standard deviations of 3 replicates (low-throughput) or propagated uncertainty from ratios of 3 experimental and 3 control measurements (high-throughput). Colored arrows correspond to example proteins shown in panel D. ✱ marks the protein rocklin_batch2_667294 because it was assayed at ∼6.5 mg/ml as higher concentrations triggered aggregation. ▲ marks the protein rocklin_batch2_104693 which appeared dimeric and not monomeric by SEC (Fig. S7). (D). Example individual protein structures and protein features compared to the distribution of metagenomic proteins (grey, both Metagenomic 1 and Metagenomic 2). ΔG_unfolding_ was measured with cDNA display proteolysis. Hydrophobicity was calculated using the scale from Monera et al. (1995), based on reversed-phase HPLC retention times of amphipathic α-helical peptides with single amino acid substitutions. pI was calculated using pK values from Bjellqvist et al. (1993 & 1994).

### Aggregation measured in mixed pools largely reflects intrinsic aggregation behavior

Because high-throughput measurements are performed in pooled experiments where co-aggregation can occur, we asked how library context influences the aggregation of single proteins. To examine this, we included 1,050 shared protein domains in both Metagenomic 1 and Metagenomic 2 libraries (roughly 10% of each library). Although the specific domains differ between the libraries, they were both drawn from the same distribution of natural domains and thus have similar feature distributions (e.g., hydrophobicity, pI, length, etc.), providing a basis for comparing aggregation behavior across independent but comparable library contexts. Across the 825 domains that were successfully quantified in both libraries, measurements under temperature stress were in close agreement, showing high correlation and low mean absolute deviation between libraries (Fig. 2B, Pearson’s r = 0.88-0.89, MAD = 0.24-0.43 log_2_(Fold Change)), consistent with the reproducibility previously reported in mixed lysate thermal proteome profiling experiments reported by Jarzab et al., 2020. Under acidic stress conditions we also observe a strong correlation, but with an offset from the y=x line reflecting a lower level of total aggregation in the Metagenomic 2 library compared to Metagenomic 1 (Fig. 2B, Pearson’s r = 0.8). This trend was consistent with the change in concentration measured for the total protein samples (Fig. S1C). We used the line of best fit to adjust the aggregation measures for all the proteins between the two libraries so they could be combined for machine learning (Fig. S3, see methods).

We next asked whether the degree of aggregation for different proteins in a mixture (where many proteins likely co-aggregate) reflects the aggregation propensity of those proteins as individually purified protein samples (each protein aggregating only with itself). To test this, we selected domains from our high-throughput dataset spanning a range of aggregation phenotypes and biophysical features for individual testing. This set also included several domains with unexpected phenotypes relative to conventional intuition—for example, low-hydrophobicity proteins that aggregated under high temperature stress, or low-pI proteins that resisted aggregation at acidic pH. We expressed and purified these domains individually (using SEC to confirm the protein is monomeric, Fig. S7) and concentrated them to the same total protein concentration of 10 mg/ml. Next, we subjected each protein to the same stress conditions and aggregation protocol used in the pooled assay (Fig. 1C), and measured the loss of soluble protein with a BCA assay. Overall, aggregation measured in a mixed pool and in individual samples correlated well (Fig. 2C, Pearson’s r = 0.66-0.78). This indicates that the aggregation measured in a pooled library generally reflects how a protein behaves on its own. Some proteins, however, were more aggregation-prone when assayed individually than in the mixed library. This could reflect greater sensitivity in the individual assays, where likely higher levels of partially or fully unfolded protein are present, or it may indicate a mechanistic preference for self-aggregation over co-aggregation with unrelated proteins. To our knowledge, this is the first time the relationship between protein aggregation in a mixed pool experiment and protein aggregation of individually pure samples has been investigated.

In addition to the overall consistency between mixed pool and single protein experiments, several counterintuitive results from our high-throughput data were also recapitulated in the low-throughput measurements. For example, the metagenomic domain rocklin_batch2_636468 (*ββββα* topology) has low hydrophobicity and high folding stability yet aggregates extensively at 75 °C (orange, Fig. 2D, ∼29% remained soluble). By contrast, the metagenomic domain rocklin_batch2_696480 (*βαββα* topology) has lower folding stability and higher hydrophobicity but is more aggregation-resistant at 75 °C (magenta, Fig. 2D, ∼92% remained soluble). Under acidic stress, the same domain rocklin_batch2_696480 aggregates despite having a high pI (green, Fig. 2D, ∼19% remained soluble), while the *de novo* designed ⍺⍺⍺ domain HHH_rd2_0006 (Rocklin et al., 2017) remains resistant despite having a low pI (blue, Fig. 2D, ∼100% remained soluble), perhaps driven by the differences in hydrophobicity (Fig. 2D). These cases highlight that the aggregation behavior of a globular protein domain following temperature or acidic stress is a complex property that cannot be simply explained by a single, one-dimensional molecular feature.

### Protein aggregation trends with protein features, but no feature explains all the variation

What protein features are important determinants of aggregation? To examine this, we predicted the structures for all the proteins using AlphaFold 2 (Jumper et al., 2021) and then computed ∼1,600 protein features derived from sequence (amino acid composition, intrinsic disorder prediction (Hu et al., 2021), hydrophobicity, etc.) and structure (Rosetta energies, exposed non-polar residues, radius of gyration, etc.) (Ferrari et al., 2025). We also measured the global folding stability of each domain using cDNA display proteolysis and included these experimental folding stability measurements in our analysis (Tsuboyama et al., 2023).

Proteins predicted to have disordered regions tend to be aggregation resistant. Aggregation resistance positively correlated with the radius of gyration of AlphaFold-predicted models, indicating that proteins with disordered regions were generally aggregation-resistant (Fig. S4A, mean log_2_(fold change) ± std: 50 °C −0.13 ± 0.46 (disordered), −0.41 ± 0.78 (compact); 75 °C −0.20 ± 0.65 (disordered), −0.69 ± 1.02 (compact); pH 4 −0.31 ± 0.51 (disordered), −0.36 ± 0.60 (compact)). This trend is driven by ∼1,800 primarily aggregation-resistant domains whose predicted structures were primarily disordered, despite substantial hydrophobicity (Fig. S4C). This is consistent with previously observed correlations between disorder and thermal stability/refoldability (Jarzab et al., 2020; To et al., 2021). Proteins not identified by MS also showed higher radii of gyration, suggesting that some aggregation-prone disordered proteins may fail to express (Fig. S4B). To focus on globular proteins, we excluded unfolded domains based on their radius of gyration (Fig. S2D), leaving 17,116 globular domains for machine learning analyses (Fig. S2A).

Across these globular proteins, aggregation propensities are only modestly related to protein topology. Within the Protein Family library, average aggregation propensity varied between families, but the trends depended on the stress applied (Fig. 3A). For example, PASTA domains aggregated more at 75 °C than at low pH, whereas SH3 domains aggregated at low pH but remained mostly soluble at 75 °C. Natural and *de novo* designed proteins aggregated to similar extents on average (Fig. 3A); although statistical comparisons were significant, the effect sizes were small, indicating that the observed differences are unlikely to be biologically meaningful (rank-biserial correlations <0.2) (Table 1). Despite the faster folding rates of ⍺-helical proteins (Kubelka et al., 2004) and the amyloid-forming propensity of β-strands (Cascio et al., 1989; Chiti et al., 2002, 2003; Sunde et al., 1997; von Bergen et al., 2001), aggregation was similar between the two classes in both the Metagenomic and Protein Family libraries (Fig. 3A). Overall, statistical comparisons again showed small effect sizes (rank-biserial correlations typically <0.1), even when p-values were significant due to large sample sizes (Table S1).

**Figure 3.**
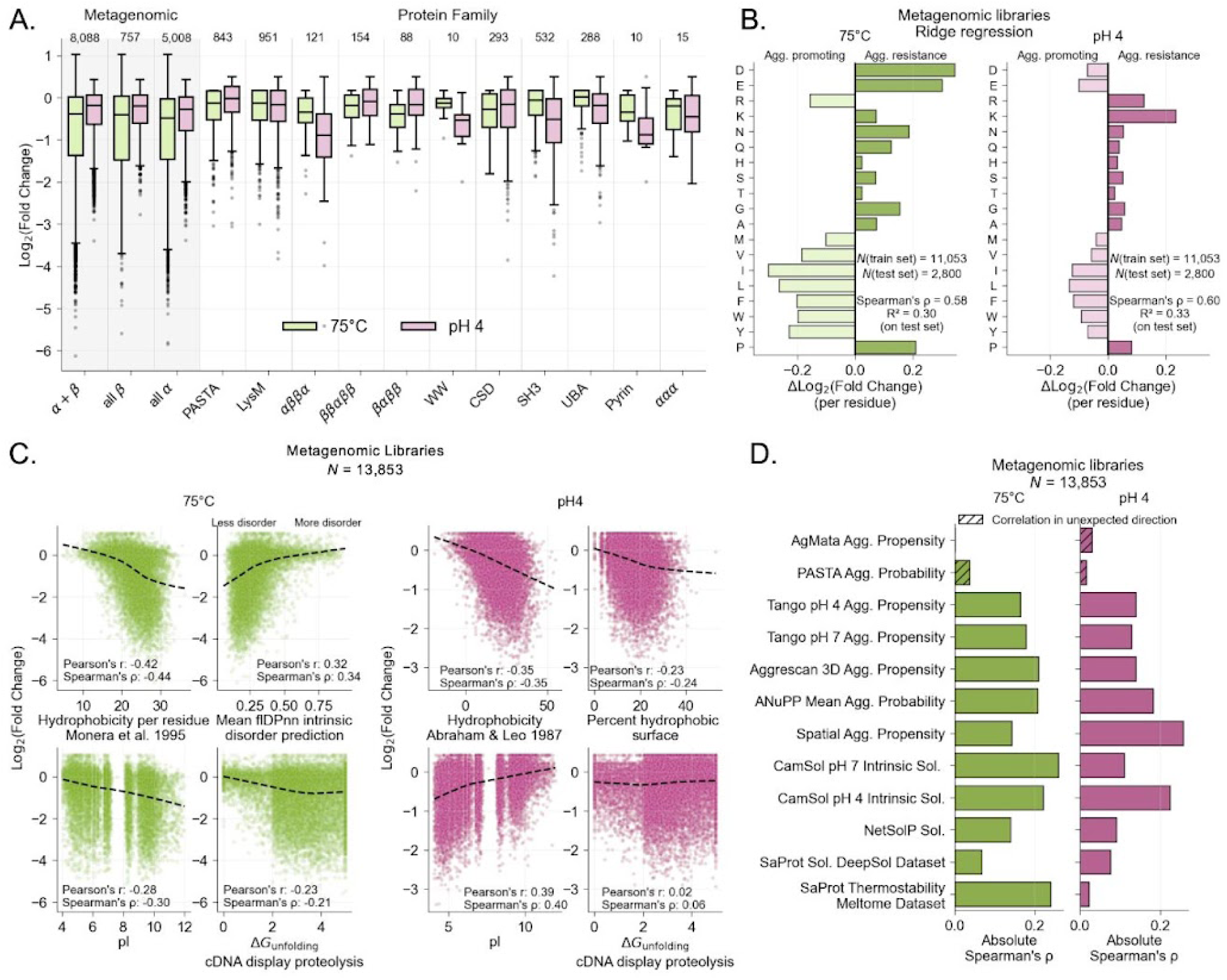
Aggregation propensity trends with protein features. (A) Distribution of heat- (75 °C, green) and acid- (pH4, pink) induced aggregation across protein topology groups in Metagenomic (left, shaded grey) and Protein Family libraries (right). Topologies for the Metagenomic libraries were determined from AlphaFold 2-predicted structures. Numbers above each box indicate the count of proteins in each topology group. (B) Ridge regression model coefficients for predicting thermal (75 °C, left, green) and acidic (pH 4, right, pink) aggregation from amino acid composition. Metagenomic sequences were clustered using MMseqs2 (Steinegger & Söding, 2017) with a minimum sequence identity of 0.3 and clusters were assigned to training and test sets to evaluate generalization. Proteins without cysteines were selected to avoid disulfide bond formation between different proteins in the mixture. (C) Aggregation trends with protein features such as hydrophobicity, intrinsic disorder (predicted by flDPnn, Hu et al., 2021) pI, ΔG_unfolding_ measured with cDNA display proteolysis. Displayed hydrophobicity scales include: Monera et al., 1995, based on reversed-phase HPLC retention times of amphipathic α-helical peptides with single amino acid substitutions; and Abraham & Leo 1987, based on fragment-based calculations of amino acid side-chain partition coefficients. (D) Absolute Spearman’s correlation between measured aggregation (75 °C: left, green; pH 4: right, pink) and published predictors of aggregation (Agg.) or solubility (Sol.). Bars with black hatching indicate directionally unexpected correlations where more aggregation prone sequences were predicted to be more aggregation resistant.

**Table 1.**
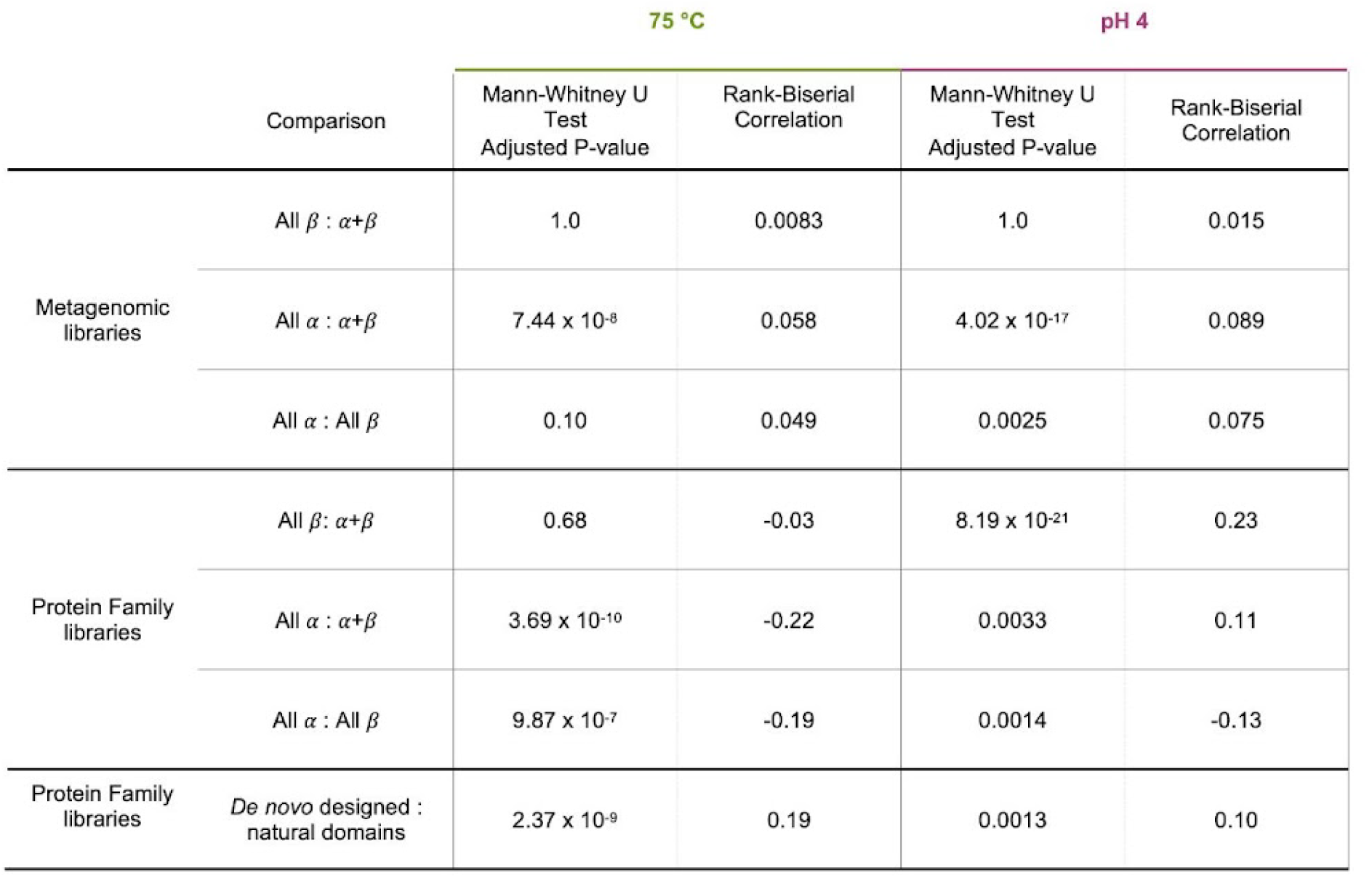
Mann–Whitney U test statistics and corresponding rank-biserial correlations for aggregation comparisons among topology groups. In libraries where multiple group comparisons were performed, p-values are adjusted using the Bonferroni correction.

|  |  | 75 °C |  | pH 4 |  |
| --- | --- | --- | --- | --- | --- |
| Comparison |  | Mann-Whitney U Test<br>Adjusted P-value | Rank-Biserial Correlation | Mann-Whitney U Test<br>Adjusted P-value | Rank-Biserial Correlation |
| Metagenomic libraries | All $\beta$ : $\alpha+\beta$ | 1.0 | 0.0083 | 1.0 | 0.015 |
| | All $\alpha$ : $\alpha+\beta$ | $7.44 \times 10^{-8}$ | 0.058 | $4.02 \times 10^{-17}$ | 0.089 |
| | All $\alpha$ : All $\beta$ | 0.10 | 0.049 | 0.0025 | 0.075 |
| Protein Family libraries | All $\beta$ : $\alpha+\beta$ | 0.68 | -0.03 | $8.19 \times 10^{-21}$ | 0.23 |
| | All $\alpha$ : $\alpha+\beta$ | $3.69 \times 10^{-10}$ | -0.22 | 0.0033 | 0.11 |
| | All $\alpha$ : All $\beta$ | $9.87 \times 10^{-7}$ | -0.19 | 0.0014 | -0.13 |
| Protein Family libraries | *De novo* designed :<br>natural domains | $2.37 \times 10^{-9}$ | 0.19 | 0.0013 | 0.10 |

**Table 2.**
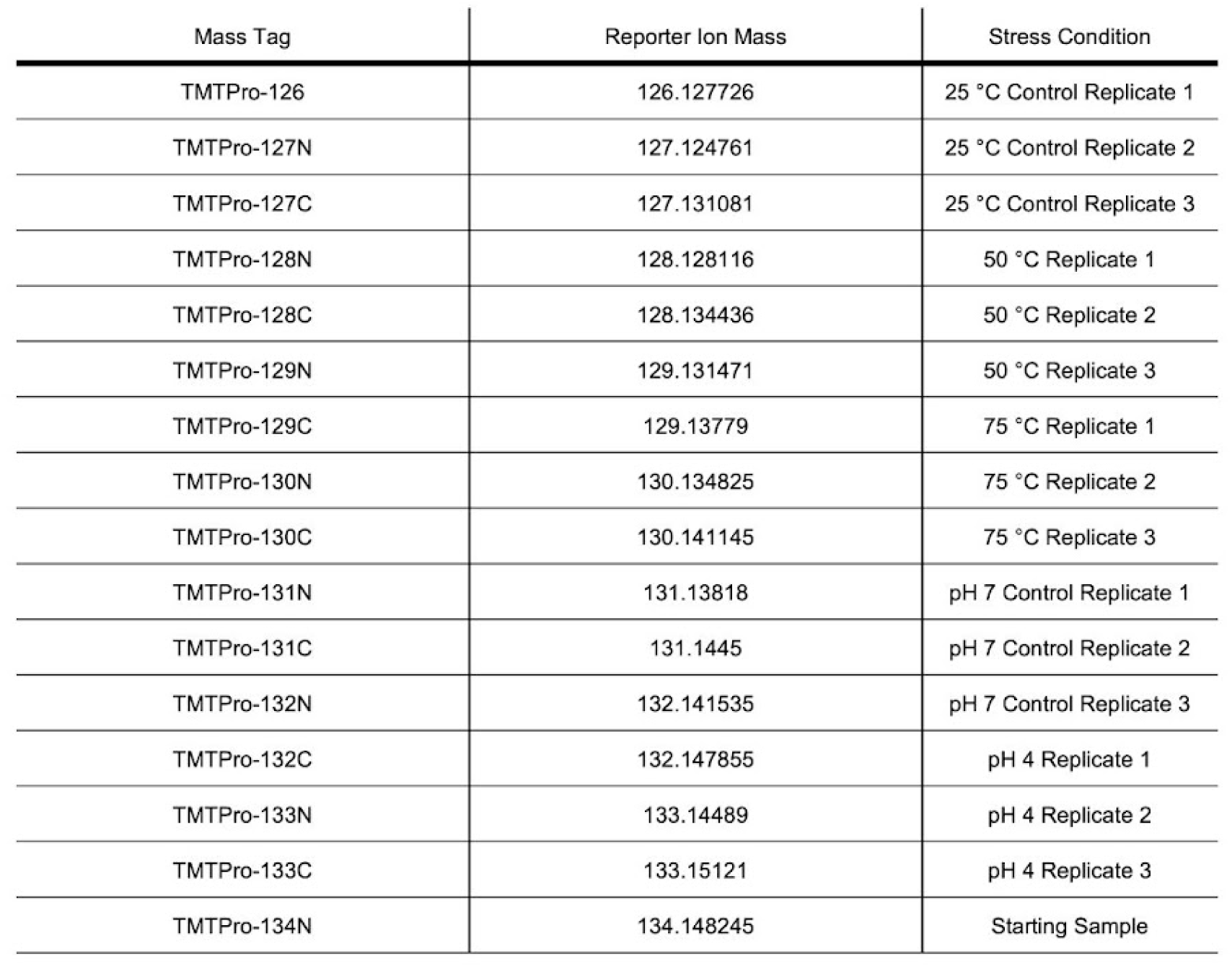
Mass tags used to label each condition. The same pairings of tags and conditions were used for all libraries.

| Mass Tag | Reporter Ion Mass | Stress Condition |
| --- | --- | --- |
| TMTPro-126 | 126.127726 | 25 °C Control Replicate 1 |
| TMTPro-127N | 127.124761 | 25 °C Control Replicate 2 |
| TMTPro-127C | 127.131081 | 25 °C Control Replicate 3 |
| TMTPro-128N | 128.128116 | 50 °C Replicate 1 |
| TMTPro-128C | 128.134436 | 50 °C Replicate 2 |
| TMTPro-129N | 129.131471 | 50 °C Replicate 3 |
| TMTPro-129C | 129.13779 | 75 °C Replicate 1 |
| TMTPro-130N | 130.134825 | 75 °C Replicate 2 |
| TMTPro-130C | 130.141145 | 75 °C Replicate 3 |
| TMTPro-131N | 131.13818 | pH 7 Control Replicate 1 |
| TMTPro-131C | 131.1445 | pH 7 Control Replicate 2 |
| TMTPro-132N | 132.141535 | pH 7 Control Replicate 3 |
| TMTPro-132C | 132.147855 | pH 4 Replicate 1 |
| TMTPro-133N | 133.14489 | pH 4 Replicate 2 |
| TMTPro-133C | 133.15121 | pH 4 Replicate 3 |
| TMTPro-134N | 134.148245 | Starting Sample |

Variation in aggregation propensity is partially explained by amino acid composition, with stress-specific effects. We focused on the Metagenomic dataset for machine learning (see methods) as it provided a broad range of aggregation behaviors under each stress and to avoid overfitting to specific folds in the Protein Family library. Using ridge regression with amino acid composition as input features, we explained 29-35% of the variation in aggregation (Fig. 3B). Hydrophobic residues generally were correlated with increased aggregation for both stresses (Fig. 3B). Acidic residues were correlated with increased aggregation at low pH, as expected because the solubilizing effect of these residues decreases in their protonated form. Acidic residues were also correlated with aggregation resistance at 75 °C, while arginine specifically increased aggregation at 75 °C (Fig. 3B, Austerberry et al., 2019; Chan et al., 2013; Ohtake et al., 2011).

Several sequence- and structure-derived features showed moderate correlations with aggregation (Fig. 3C, Fig. S8B-D). Many features were correlated with each other, so we clustered them and analyzed the most strongly correlated feature in each cluster (Fig. S8A). At both 50 °C and 75 °C, similar features emerged as important, including hydrophobicity, pI, predicted intrinsic disorder, and Rosetta energetic terms, all showing correlations in the same direction for both stresses (Fig. 3C, Fig. S8B,C; |Pearson’s r| > 0.2). Surprisingly, increased folding stability (experimentally measured by cDNA proteolysis, Tsuboyama et al., 2023) correlated with increased aggregation at 75 °C, but was not a strong feature for 50 °C aggregation (Fig.3C; 75 °C Pearson’s r = −0.23). This result could be confounded by other protein features, like hydrophobicity, that simultaneously promote high stability and aggregation (Chiti et al., 2002; Dill, 1990; Santos et al., 2020, and see Fig. 4 discussion below). For acidic stress, we see strong correlations with hydrophobicity, surface hydrophobics, composition of negatively charged residues, net charge, and pI (Fig. 3C, Fig. S8D |Pearson’s r| = 0.19-0.39). Unlike the 75 °C stress, there was little correlation between pH-induced aggregation and experimentally measured folding stability at pH 7 (Fig. 3C).

**Figure 4.**
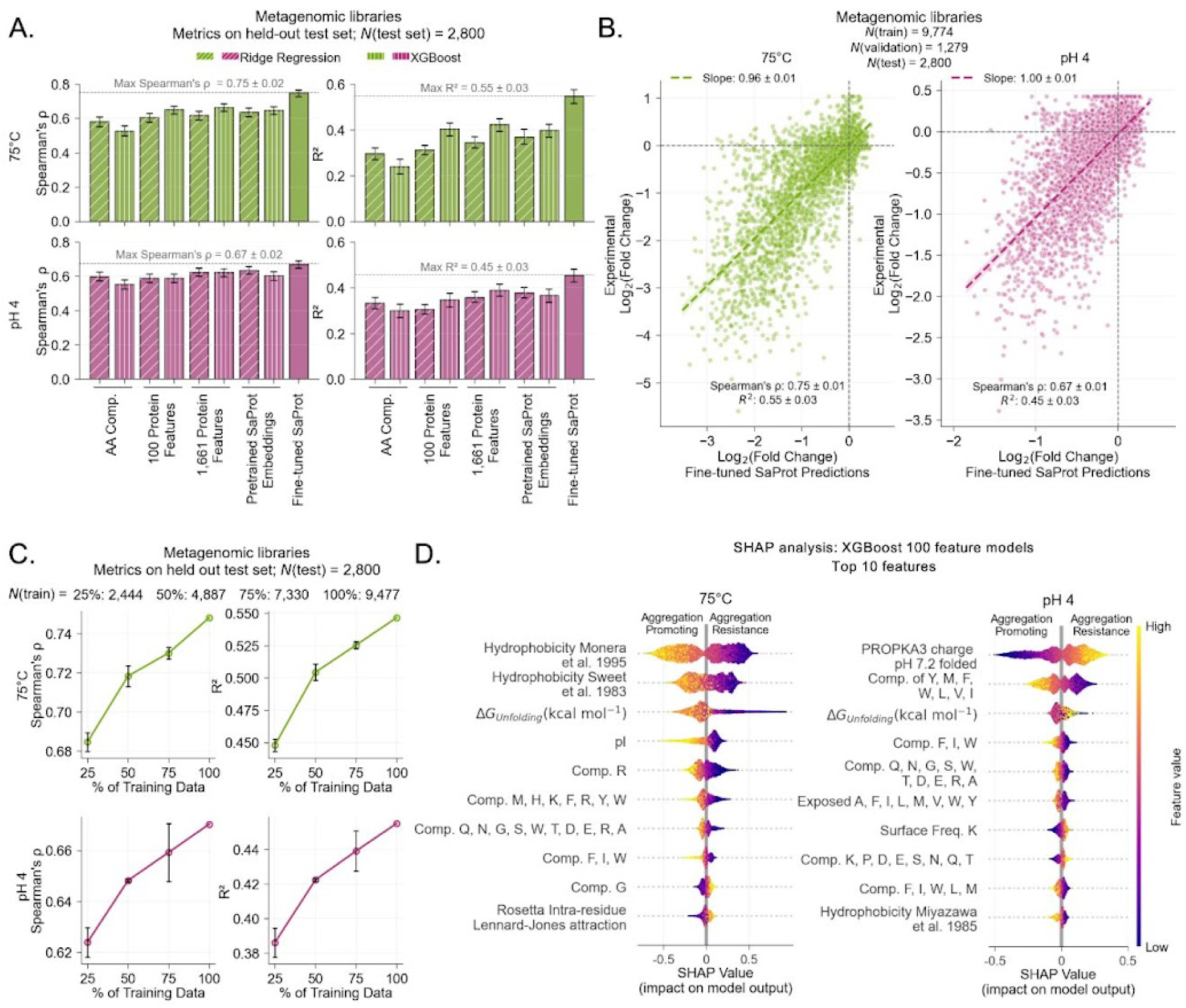
Fine-tuned SaProt pretrained language models predict stress-induced aggregation. (A) Predictive performance of different model architectures and input features for thermal (75 °C, top, green) and acidic (pH 4, bottom, pink) aggregation. Metrics were evaluated on a held-out test set using MMseqs2 clustering as described in figure 3 legend and the methods. Error bars represent half the 95% CI from bootstrapping. Bar patterns indicate the model architecture: diagonal hatching for ridge regression, vertical hatching for XGBoost, solid fine-tuned language models. (B) Comparison of experimental and predicted aggregation values from best-performing models (fine-tuned SaProt models: 75 °C, left/green; pH 4, right/pink) on the held-out test set. Statistics are reported as mean ± half 95% CI from bootstrapping. (C) Performance of fine-tuned SaProt models on the test set when trained on subsets of the training data. The points and error bars represent the mean and standard deviation across three models trained on random subsets of the train set. (D) SHAP analysis of XGBoost model trained on 100 features, showing feature importance for predicting heat- (75°C, left) and acid- (pH 4, right) induced aggregation. Each point is colored by the value of the feature where higher values are more yellow and lower values are more purple.

Overall, individual features explain only modest variation. Even the strongest correlated features like hydrophobicity or pI fail to explain the majority of variation in aggregation (Fig. 3C). For example, there are highly hydrophobic proteins that don’t aggregate at 75 °C and there are low pI proteins that don’t aggregate at low pH (Fig. 3C). This indicates the complexity of aggregation and the need for models that consider the contribution and interaction between many protein properties.

### Published aggregation and solubility prediction tools show limited generalization to our data

Existing predictors of aggregation (Chennamsetty et al., 2009; Conchillo-Solé et al., 2007; Fernandez-Escamilla et al., 2004; Orlando et al., 2020; Prabakaran et al., 2021; Trovato et al., 2007), solubility (Sormanni et al., 2015; Su et al., 2025; Thumuluri et al., 2022), and thermostability (Su et al., 2025) showed a range of relatively weak correlations with our measurements, reflecting both the limitations of existing models and differences in aggregation mechanisms being modeled (Fig. 3D). Many of these tools were developed using data on amyloid-β peptides or on aggregation-related phenotypes measured for globular proteins. Because they were not trained to predict aggregation under the forced stress conditions used in our experiments, perfect agreement with our data is not expected. Consistent with this expectation, published models show weak correlations with our measured aggregation, with absolute Spearman’s correlations ranging from 0 - 0.26. CamSol (Sormanni et al., 2015) was the strongest model for 75 °C aggregation (Fig. 3D, |Spearman’s ⍴| = 0.26) and spatial aggregation propensity (SAP, Chennamsetty et al., 2009) calculated from the static predicted structure performed best for acid-induced aggregation (Fig. 3D, |Spearman’s ⍴| = 0.26). Interestingly, SAP was a stronger predictor of pH 4-induced aggregation (Fig. 3D, |Spearman’s ⍴| = 0.26) than it was of aggregation after 75 °C (|Spearman’s ⍴| = 0.14). Because SAP is computed based on hydrophobic patches on a protein’s putative native structure, this suggests aggregation at pH 4 is more likely to be driven by the native folded state compared with aggregation from 75 °C stress. CamSol pH 4 was a better predictor of acid-induced aggregation than CamSol pH 7, indicating that prediction accuracy improves when the modeled pH matches the experimental pH. Additionally, the SaProt language model finetuned on the Meltome dataset (measurements of apparent (non-equilibrium) melting temperature for the human proteome in the cell, Jarzab et al., 2020; Su et al., 2025) was nearly the most accurate model predicting 75 °C aggregation, but not acid-induced aggregation. Still, the overall accuracy of the Meltome trained model was low (Fig. 3D, |Spearman’s ⍴| = 0.24). These results motivated us to develop models specifically tailored to predict aggregation under the stresses we measured.

### Fine-tuned SaProt language models generalize well to predict protein aggregation

To enable future engineering of aggregation-resistant proteins and prioritization of candidates for experimental testing, we built predictive models of aggregation resistance from the Metagenomic dataset using protein sequence and structure features. We evaluated three machine learning strategies that balance interpretability and predictive power: ridge regression, XGBoost, and fine-tuning SaProt, a language model trained on structurally aware sequences generated with FoldSeek (Su et al., 2024, 2025; van Kempen et al., 2024). To test generalization and avoid overfitting, we clustered sequences using MMseqs2 (Steinegger & Söding, 2017) at 30% sequence identity (Fig. S9A) and split the clusters into train-test-validation sets (Fig. S9B-C, the mean pairwise sequence identity between sets is 23%), ensuring similar distributions of aggregation phenotypes and protein features across sets (Fig. S9D-E).

Overall, our fine-tuned SaProt models outperformed all other approaches in predicting protein aggregation for both aggregation stresses (Fig. 4A). Fine-tuning SaProt models explained 45% of the variance in acid-induced aggregation, 43% in 50 °C-induced aggregation and 55% in 75 °C-induced aggregation (Fig. 4A, Fig. S11). Overall, the model generalized well on the held-out test set, although it underestimated the degree of aggregation for some proteins (Fig. 4B). Structural information was important for prediction accuracy and using structures from Boltz-2 (Passaro et al., 2025) and AlphaFold 2 (Jumper et al., 2021) for test set predictions showed similar performance (Fig. S12). Acid-induced aggregation predictions were less dependent on structural input than 75 °C aggregation predictions (Fig. S12). Subsampling the training set and evaluating on the held-out test set showed that model performance continues to improve with more data, indicating collecting more data could further improve predictions (Fig. 4C). In interpretable models, prediction accuracy improved with the inclusion of more protein features, although gains were modest between 100 to 1,661 features (Fig. 4A, Fig. S10). Models incorporating sequence- and structure-based inputs outperformed those using amino acid composition alone (Fig. 4A).

Because the XGBoost model provided greater interpretability than the SaProt model while only incrementally sacrificing accuracy (Fig. 4A), we used it to examine how protein features contribute to aggregation predictions using Shapley Additive Explanations (SHAP) (Lundberg et al., 2020; Lundberg & Lee, 2017). For each prediction, SHAP analysis quantifies how much each feature contributes by representing the model output as a sum of locally additive feature effects, even when the underlying model is a nonadditive model. Figure 4D summarizes the combined results across all predictions. The most important feature for predicting aggregation at 75 °C was hydrophobicity calculated from the primary sequence using a scale based on chromatographic measurements of point mutated peptides (Monera et al., 1995). Net charge at pH 7.2, calculated with PROPKA3 (Olsson et al., 2011) from the predicted AlphaFold structure, was the most predictive for acid-induced aggregation (Fig. 4D).

The top ten features were primarily sequence-based measures of hydrophobicity and amino acid composition (Fig. 4D). Folding stability measured by cDNA display proteolysis (Tsuboyama et al., 2023) also emerged as a top feature for predicting both stress conditions. Notably, even after accounting for other features, low folding stability contributed to aggregation resistance following 75 °C stress (Fig. S13A). For acid-induced aggregation, SHAP analysis found that intermediate folding stability (∼1.5 kcal/mol < ΔG_unfolding_ < ∼3.5 kcal/mol) contributed to increased aggregation, whereas higher or lower stability contributed to aggregation resistance, although these contributions were approximately 3-4 fold smaller than for aggregation following 75 °C stress (Fig. 4D, Fig. S13A,B). The predominance of sequence-based features among the top predictors is consistent with the aggregation we observe occurring from partially folded or unfolded states rather than the fully folded structure, but may also indicate we lack appropriate descriptors for how the folded structure relates to aggregation. Future predictive models may improve by incorporating experimental or computational measures of protein dynamics and features derived from these alternative conformations.

### SaProt 75 °C predictions show weak but consistent trends across related experimental datasets

Although our models were trained to predict heat- and acid-induced aggregation of small protein domains, we investigated whether our models are useful for out-of-domain predictions for related phenotypes. We compared predictions from our SaProt 75 °C model to experimental datasets of soluble expression (Niwa et al., 2009; Uemura et al., 2018), thermostability (Jarzab et al., 2020), mutation effects on solubility (Attanasio et al., 2025), and amyloid properties of peptides (Louros et al., 2020; M. Thompson et al., 2025). Across all datasets, correlations between predicted aggregation resistance and experimental measures of solubility, apparent melting temperature, or solubility of mutants were positive, but weak (Pearson’s r ≅ 0.1-0.3, Fig. S14A-D). For amyloid peptide datasets, non-amyloid peptides were predicted to be slightly more aggregation resistant than amyloid-β peptides, and non-nucleating sequences were predicted to be more aggregation-resistant than nucleating sequences. (Fig. S14F). Incorporating SaProt predictions with amino acid composition in regression models to predict these properties showed limited or no improvement over just amino acid composition alone (Fig. S14G). Overall, these results indicate that the SaProt 75 °C model shows limited predictive power for these external datasets, although it weakly captures general trends related to protein solubility and aggregation that extend beyond its original training domain and protein size range.

### SaProt predictions of single mutants reflect experimentally observed trends

Protein engineers commonly seek to reduce aggregation propensity while modifying a protein’s sequence with as few substitutions as possible. Can our fine-tuned SaProt model predict the effects of single substitutions even though it was trained on protein domains with low sequence identity to each other? This is difficult to evaluate experimentally at scale due to the limitations of bottom-up mass spectrometry. Since direct testing of thousands of single mutants is impractical, we leveraged natural variation in clusters of related proteins in our dataset as an indirect method to estimate the effects of single substitutions on the aggregation phenotypes we observed.

To estimate experimental effects of amino acid substitutions within our held-out test set, we analyzed 61 sequence clusters (≥5 proteins each) containing 1,480 aligned positions with sufficient amino acid diversity and representation across amino acid groups (see methods); an example cluster is shown in Fig. 5A-F. For each position, we grouped together sequences with related amino acid identities at that position (Fig. 5G, Fig. S15C, see methods for how amino acids were grouped) and averaged our aggregation measurements (log_2_ fold change following 75 °C stress) for the sequences in the group to estimate an overall aggregation propensity for each amino group at a given position (Fig. 5G). Finally, we estimated the effects of amino acid substitutions from one group to another by taking the difference in the average aggregation measurements of sequences in each group. For example, within cluster 105 (Fig. 5B), the average fold-change for sequences with Ile, Leu, or Tyr (group ILY) at position 27 is −2.1, and the average fold-change for sequences with Asp or Glu (group DE) at position 27 is −0.5 (Fig. 5G). We estimated the experimental effect of substituting position 27 from group DE to group ILY as the difference ILY - DE = −1.6, meaning sequences with D or E at position 27 aggregate about 3-fold less than those with I, L, or Y. It is important to note that this estimate is very approximate, because the sequences in cluster 105 have pairwise sequence identities less than 51%, and the differences in aggregation between these groups cannot fully be attributed to sequence variation at position 27. For each aligned position, we compared mean aggregation between amino acid groups, but site-specific differences were not statistically significant after correcting for multiple testing. However, aggregating these results across all 1,480 aligned positions enables us to indirectly evaluate our fine-tuned SaProt model at predicting the trends observed between amino acid groups at individual positions in a large-scale, statistically robust way.

**Figure 5.**
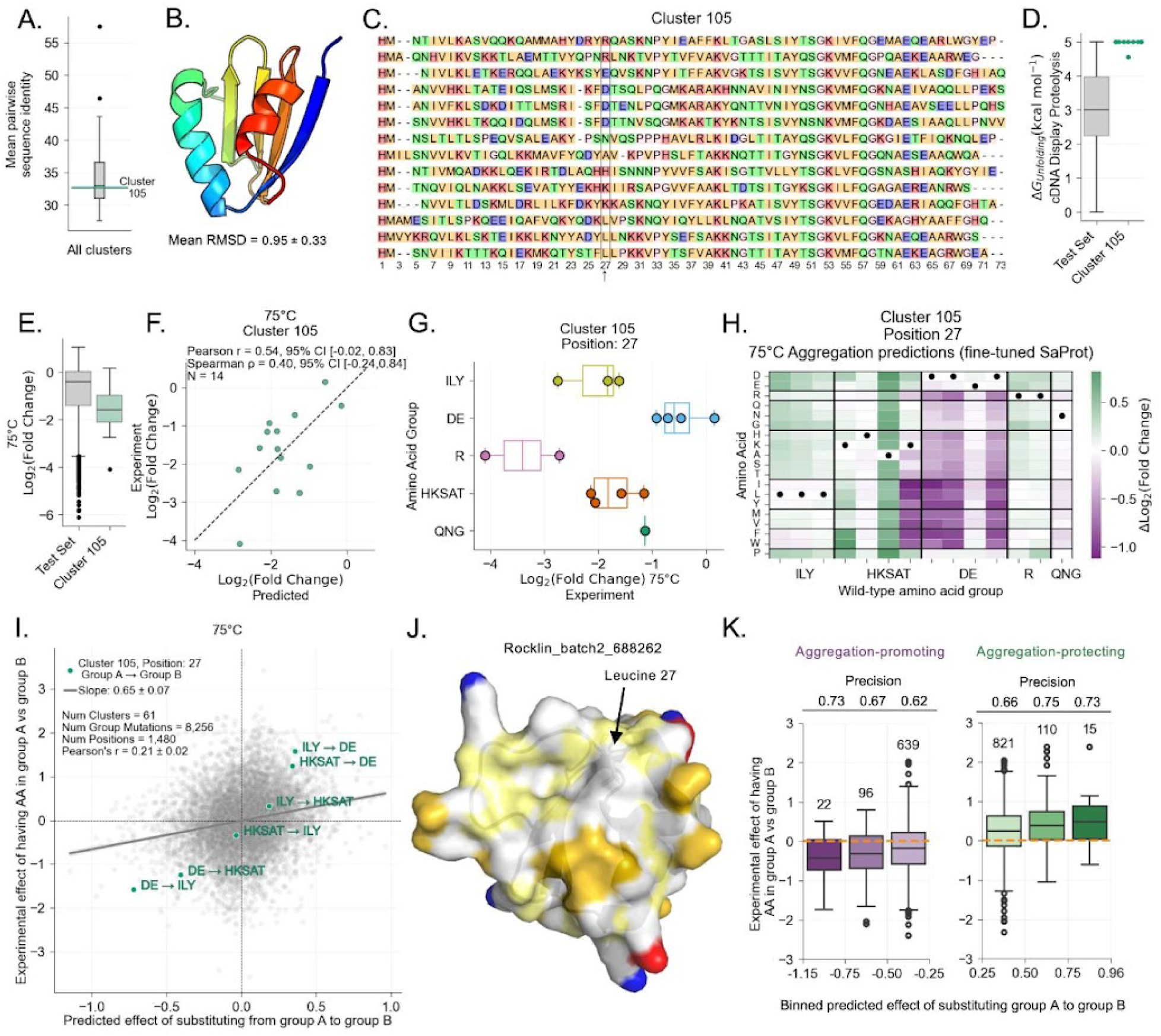
Deep mutational scanning predictions correlate with experimental trends. (A) Distribution of mean pairwise sequence identity within each cluster in the test set. The mean for cluster 105 is highlighted in teal. (B) Example topology of cluster 105. The AlphaFold 2-predicted structure for rocklin_batch2_202839 is shown, along with the mean and standard deviation RMSD calculated between all pairs of structures in the cluster. (C) Sequence alignment of the 14 proteins in cluster 105, colored by amino acid properties. (D) Distribution of experimental ΔG_unfolding_ measured by cDNA display proteolysis for all proteins in the test set (grey) and cluster 105 (teal) (E) Distribution of experimental log_2_(fold change) at 75 °C for all proteins in the test set (grey) and cluster 105 (teal). (F) Comparison of experimental and fine-tuned SaProt predicted log_2_(fold change) at 75 °C for the wild-type proteins in cluster 105. (G) Distribution of experimental log_2_(fold change) at 75 °C for wild-type proteins with amino acids in different groups at position 27. (H) Heatmap of Δlog_2_(fold change) (fine-tuned SaProt prediction minus wild type prediction) for all single point mutations at position 27. Positive values (green) indicate mutations predicted to increase aggregation resistance while negative values (purple) indicate the mutation is predicted to increase aggregation propensity. Wild-type residues at position 27 are marked with a black dot and proteins are grouped by the amino acid group of the wild-type residue. (I) Comparison of experimental differences between wild types with group A vs group B at a given position and fine-tuned SaProt predicted effects for substitutions from group A to group B. Analyses were performed within clusters for all test set clusters with ≥ 5 proteins. Slope and Pearson’s r are shown as mean ± half 95% CI from bootstrapping. Teal points highlight comparisons for position 27 in cluster 105. (J) Example structure of rocklin_batch2_688262, a protein in cluster 105 with a leucine at position 27. Colored by properties of the residue (red: negative, Asp/Glu side chain; blue: positive, Arg/Lys side chain; dark yellow: polar side chains, Ser/Thr/Asn/Gln/His/Trp/Tyr; light yellow: surface backbone atoms, C/O/N; white: hydrophobic side chains). (K) Distribution of experimental effect between wild types with group A vs group B at a given position for the stronger SaProt predictions for substitutions from group A to group B. Boxes are labelled with the number of proteins in each bin and the precision (calculated by the ratio of true positives to all predicted positives). Predictions are split into aggregation-promoting substitutions (purple, left) and aggregation-protecting substitutions (green, right). The orange dotted line indicates y = 0.

We used our 75 °C SaProt model to predict the change in aggregation for all possible single substitutions for each protein in the test set (Δlog_2_(fold change), Fig. 5H). Most substitutions had small predicted effects, although a subset showed large predicted changes in aggregation (Fig. S15A). To summarize the predicted effect of substituting from a wild-type amino acid group (e.g. all proteins in cluster 105 with I, L, or Y at position 27) to a target amino acid group (eg. group DE), we averaged the predicted Δlog_2_(fold change) for all substitutions at the given site to all target amino acids (Fig. 5H). For example, in cluster 105, the average Δlog_2_(fold change) for predicted substitutions at position 27 from group ILY (wild-type amino acid) to group DE was +0.72, indicating these substitutions are predicted to reduce aggregation (Fig. 5I).

These predicted effects showed a significant correlation with the experimentally observed trends (Fig. 5I). Across all 1,480 positions in 61 clusters, we observed a positive correlation and slope between predicted and experimental effects of single substitutions (Fig. 5I, Pearson’s r = 0.21 ± 0.02, slope = 0.65 ± 0.06, mean ± half the 95% CI computed from bootstrapping). Restricting the analysis to positions and amino acid groups with more proteins—thereby reducing the dataset size—showed similar trends (Fig. S15E). To illustrate, at position 27 in cluster 105, predicted substitutions from group DE to group ILY were the most aggregation-promoting (Δlog_2_(fold change) = −0.72), consistent with the largest experimental difference observed between wild-type proteins with DE versus ILY at this site (Δlog_2_(fold change) = −1.6). Substitutions from HKSAT to ILY were predicted to have little impact, consistent with the estimated experimental phenotypes (Fig. 5I, teal). Position 27 lies in a loop near the protein surface, where a leucine at this position creates a hydrophobic patch on the surface (Fig. 5J). Importantly, the strongest predicted effects were generally consistent with experimentally observed trends (Fig. 5K). We quantified this consistency using precision, defined as the fraction of the strongest predictions (all positives) whose direction matched the experimentally observed trends (true positives). For the strongest predictions (absolute predicted effect > 0.5), we see 71% of the predicted effects matched the experimentally observed direction of substitution effects (precision > 0.71, Fig. 5J), with experimental differences typically 1.3 times as large as predicted effects. Many experimentally observed effects were not predicted, but these could stem from coincidental (not causative) effects due to the large sequence difference between the proteins in these clusters. Along with the overall accuracy of the predictive model (Fig. 4A,B), the consistency between the predicted substitutions and experimentally observed trends at individual sites (Fig. 5I), especially for the strongest predictions (Fig. 5K), highlights the potential for applying our model in protein engineering.

## Discussion

Understanding and predicting protein aggregation remains a challenge in drug development and biotechnology. Here, we quantified a range of aggregation phenotypes across thousands of small protein domains (Fig. 1D). By using custom-designed libraries of model proteins, we were able to compare related and dissimilar structures to quantify how topology, sequence composition, and structural features influence aggregation behavior. Additionally, our in vitro approach avoided the complexities of cellular proteomes and enabled direct comparison of aggregation induced by different stresses, revealing that aggregation behavior was stress dependent (Fig. 1E). Importantly, aggregation measurements from pooled assays correlated with results from individually tested proteins (Fig. 2C), indicating that aggregation is strongly influenced by intrinsic protein properties as well as environmental context. This finding supports the use of multiplexed, high-throughput approaches to measure aggregation without purifying each protein individually. Feature-based analyses identified several properties that correlated with aggregation propensity, but no single feature or combination of features fully explained the observed variation (Fig. 3), underscoring the complexity of predicting aggregation in globular proteins. Fine-tuning the SaProt language model yielded the most accurate aggregation predictions, explaining 43-55% of the variance in a held-out test set (Fig. 4A), and incorporating structural information was important for predictive performance (Fig. S12).

Our approach has several limitations. First, because we quantified changes in relative protein abundance due to insoluble aggregation, our measures of aggregation resistance do not guarantee that “non-aggregating” proteins remain monomeric and folded in their native structures. Second, our assay does not distinguish among the different mechanisms of aggregation, which can complicate modeling. Third, our approach is limited to proteins that express solubly, which could bias the sequence space analyzed in this assay. Still, this bias appears minimal, as the identified proteins spanned a wide range of protein features and covered ∼82% of the ordered domains (Fig. S2). Our computational analyses also indicate that machine learning performance for predicting protein aggregation has not yet plateaued (Fig. 4C). Integrating large biophysical datasets on protein conformational dynamics (Ferrari et al., 2025) could further improve modeling, which currently relies on predicted native folded-state structures and sequence features, even though aggregation may arise from partially unfolded intermediates.

Collectively, our work provides a large, valuable dataset of high temperature- and acidic-stress induced insoluble aggregation across thousands of diverse globular protein domains, establishing a foundation for understanding and predicting protein aggregation. Our approach can be expanded to address key unanswered questions in the field, including measuring aggregation driven by diverse mechanisms under different stress conditions, determining how stress-induced aggregation relates to long-term stability, and performing large-scale studies of how different excipients modulate aggregation in a protein-specific manner. Additionally, our approach can be applied iteratively to design and test aggregation-resistant proteins, enabling cycles of computational design and large-scale experimental validation using oligo pools. With continuing advances in DNA synthesis and mass spectrometry, our approach can generate larger, more diverse aggregation datasets to accelerate the development of predictive models and the design of aggregation-resistant proteins.

## Methods

### Library construction

#### Protein Family Library

We selected natural domains from the Pfam database (Mistry et al., 2021) and *de novo* designed small protein domains (Kim et al., 2022); (Rocklin et al., 2017). In total we selected 3,763 domains from the following *de novo* designed topologies: *βαββ, ββαββ, αββα, ααα* and natural topologies: Pyrin, Ubiquitin Binding Associated (UBA), SH3, WW, Cold Shock Domain (CSD), PASTA, LysM.

#### Metagenomic Libraries

We selected small protein domains from the MGnify database (Richardson et al., 2023) to increase the diversity of the domains studied. We split these domains into two libraries termed Metagenomic 1 and Metagenomic 2. To investigate the role of library context, one thousand metagenomic domains were selected to be present in both libraries. We also included 50 proteins that were assayed in the protein family library. In total, we selected 10,050 domains in the Metagenomic 1 library and 10,505 domains in the Metagenomic 2 library.

For all libraries, proteins were selected to have at least 2 unique tryptic peptides between 7 and 35 amino acids in length (Fig. 1C) to maximize the ability to identify the proteins with bottom-up mass spectrometry. Additionally, we excluded proteins with cysteine residues to avoid disulfide bond formation between different proteins in the library.

#### DNA ordering

The selected amino acid sequences were reverse translated and codon-optimized based on *E. coli* codon frequencies using DNAWorks2.0 (Hoover & Lubkowski, 2002). Libraries were then purchased as oligo pools from Twist Biosciences. Unique adapter sequences for each library were added to the encoding DNA sequences to enable amplification of the individual library and to add NdeI and XhoI cut sites.

### Library cloning and expression

The ordered oligonucleotide library was resuspended and amplified using SsoAdvanced Universal SYBR green Supermix (BioRad). The number of cycles for amplification to 50% the maximum signal was determined from a test qPCR run to avoid overamplification. Amplified inserts and a custom backbone, pGR02, were digested with restriction enzymes, NdeI and XhoI (NEB), and ligated using T4 DNA ligase (Thermofisher). The ligation mixture was transformed into electrocompetent *E. coli* cells (NEB) and plated on MDAG-11 non-inducing minimal media agar plates with kanamycin. All transformants were pooled to obtain transformants for the whole library and cloned plasmids were isolated using the QIAprep Spin Miniprep kit (Qiagen).

The cloned library plasmids were transformed into electrocompetent BL21(DE3) cells (Sigma-Aldrich) and plated on MDAG-11 non-inducing minimal media agar plates with kanamycin overnight at 37 °C. All transformants were pooled and used to inoculate bulk cultures of Miller LB Broth with 50 µg/ml kanamycin. Cells were grown at 37 °C, 220 rpm until OD_600_ ∼0.6. Expression was induced with a final concentration of 500 mM Isopropyl β-D-1-thiogalactopyranoside (IPTG) and cells were grown overnight (∼16-18 hrs) at 16 °C, 220 rpm. Cells were harvested by centrifugation at 4 °C.

### Individual protein cloning and expression

We selected protein domains from our high-throughput dataset to test aggregation in a low-throughput approach. We ordered gene fragments from twist biosciences encoding the selected proteins and cloned them into the pGR02 vector using Nde1 and Xho1 restriction enzyme digest and T4 DNA ligase ligation. The plasmids were transformed into BL21(DE3) chemical competent cells and grown at 37 °C, 220 rpm until OD_600_ ∼0.6. Expression was induced with a final concentration of 500 mM Isopropyl β-D-1-thiogalactopyranoside (IPTG) and cells were grown overnight (∼16-18 hrs) at 16 °C, 220 rpm. Cells were harvested by centrifugation at 4 °C.

### Library and individual protein purification

Harvested cells were lysed with lysis buffer (500 mM NaCl, 30 mM Imidazole, 0.25% CHAPS, 20 mM Tris-HCl, pH 8) and sonication. The lysate was clarified by centrifugation (13,000 x g, 45 min) and purified using immobilized metal affinity chromatography with Ni-NTA beads (QIAGEN). Beads were washed (500 mM NaCl, 30 mM Imidazole, 0.25% CHAPS, 5% Glycerol, 20 mM Tris-HCl pH8) and then protein was eluted (300 mM NaCl, 500 mM Imidazole, 5% Glycerol, 20mM Tris-HCl pH 8). Proteins were dialyzed (300 mM NaCl, 5% Glycerol, 20 mM Tris-HCl pH 8) to remove imidazole and the N-terminal His-SUMO tag was cut using ULP1 (produced in-house, Addgene plasmid #113671, pCDB327, (Lau et al., 2018)) (1:100 molar excess, overnight 4 °C). The cut SUMO tag and ULP1 were removed with another IMAC Ni-NTA purification. Proteins were further purified using size exclusion chromatography (Superdex 75 increase 10/300 GL, GE Healthcare) on a Bio-Rad NGC Chromatography System, eluting in PBS pH 7.4. We used the SEC trace collected on our instrument for a standard set of proteins (BSA: 67 kDa, Ovalbumin: 43 kDa, Ribonuclease A: 13.7 kDa, Aprotinin: 6.5 kDa, Vitamin B12: 1.4 kDa) to calculate expected volumes of elution based on molecular weight (Fig. S1B). Protein size and appearance was confirmed using SDS-PAGE and fractions containing the monomeric protein were pooled and concentrated (Amicon Ultracel-4, Millipore Sigma). Protein concentration was determined using Bicinchoninic acid (BCA) assay (ThermoScientific).

### High Throughput Aggregation assay

The purified protein library was concentrated to 10 mg/mL (Amicon Ultracel-4, Millipore Sigma), quantified by BCA Assay. The library was then aliquoted for 3 replicas of each control and stress condition. For high temperature stress, protein samples were incubated at 50 °C or 75 °C in a PCR thermocycler for 20 min and control samples were held at room temperature for 20 min. Samples were cooled on ice for 10 min, and aggregates were removed by centrifugation at 21,100 x g for 15 min. For acidic stress, the pH of the protein library sample was adjusted by adding citrate-phosphate buffer (pH 2.2 for the pH 4 stress or pH 7.4 for the control) at a 1:5 buffer-to-protein sample ratio, resulting in a pH 4 for the stress sample (based on the protocol from Oyetayo & Kiefer, 2016). The ionic strength of both citrate-phosphate buffers was adjusted with NaCl so both buffers had the same ionic strength (0.5 M) (Elving et al., 1956). Samples were incubated with shaking for 10 min, and aggregates were removed by centrifugation at 21,100 x g for 15 min. The supernatants from each sample were then collected and processed for mass spectrometry analysis.

### Low Throughput Aggregation assay

We selected individual proteins representing a range of aggregation phenotypes and protein features to test aggregation at high temperature and low pH in low-throughput. Purified proteins were concentrated to 10 mg/mL (Amicon Ultracel-4, Millipore Sigma), quantified by BCA Assay, except for rocklin_batch2_667294, which was assayed at 6.5 mg/mL because higher concentrations caused spontaneous aggregation. For high temperature stress, samples were incubated at 50 °C or 75 °C in a PCR thermocycler for 20 min, while control samples were held at room temperature for 20 min. Samples were then cooled on ice for 10 min, and aggregates were removed by centrifugation at 21,100 x g for 15 min. The protein rocklin_batch2_5453 formed a “jelly-like” pellet which was removed by filtration. For acidic stress, the pH was adjusted by adding citrate-phosphate buffer (pH 2.2 for the pH 4 stress or pH 7.4 for the control) at a buffer-to-protein ratio determined individually for each protein, resulting in a pH of 4 for the stress sample (based on the protocol from Oyetayo & Kiefer, 2016). The ionic strength of both citrate-phosphate buffers were adjusted with NaCl to have the same ionic strength (0.5 M) (Elving et al., 1956). Samples were incubated with shaking for 10 min, and aggregates were removed by centrifugation at 21,100 x g for 15 min. The supernatants were then quantified by BCA assay. Aggregation propensity was calculated as the log_2_ ratio of control sample to stress sample concentrations. For each protein, the aggregation assay was done in triplicate. The protein rocklin_batch2_104693 appeared dimeric by SEC but was nonetheless measured for aggregation and is denoted in Figure 2C.

### cDNA display proteolysis

Protein-cDNA complexes were prepared and stability was measured with proteolysis followed by next generation sequencing using the protocol described by Tsuboyama et al. 2023.

### Mass spectrometry sample preparation

The supernatants from the aggregation experiment were incubated with 7.5 M guanidine hydrochloride, 100 mM triethylammonium bicarbonate (TEAB) (final guanidinium hydrochloride concentration of 6 M) for 4 min at 95 °C to denature the proteins. We then added DTT at a concentration of 5 mM and incubated at room temperature for 20 min. We added Iodoacetamide to 15mM and incubated at room temperature for 15 min, in the dark, alkylation was then quenched with DTT. We then diluted the sample with 100mM TEAB to decrease the guanidine hydrochloride concentration below 1.5 M. Then samples were digested with 1:100 MS-grade trypsin (ThermoScientific) to sample ratio and incubated overnight at 37 °C and quenched at −80 °C. Samples were then cleaned up with C18 HyperSep columns (ThermoScientific) and the eluted sample was dried to save for further processing.

We resuspended the dried samples in 100 µl of 100 mM HEPES (pH 8.5) and incubated with shaking for 30 minutes at room temperature until fully resuspended. We then performed a microBCA assay (ThermoScientific) to determine the peptide concentration. For downstream analysis, we transferred 75 µg of peptides into a new tube and adjusted the volume 80 µl with 100 mM HEPES pH 8.5. To aid in normalization of MS signals, we used the same peptide amount for each condition. Each sample was then labelled with unique TMTPro 16-plex tags (ThermoScientific, Cat# A44520) (Table S2). We incubated the TMTPro 16-plex tags to room temperature and added the samples to the tags and incubated at room temperature for 2 hours. Labelling was quenched with 2 ul of 50% hydroxylamine and followed by a 15 minute incubation with shaking at room temperature. The labelled peptides were then fractionated with high pH reverse-phase fractionation (ThermoScientific Cat #84868) and fractions were dried prior to MS analysis.

### TMT-muliplexed tandem mass spectrometry

Three micrograms of each fraction were resuspended in buffer A (contained 94.785% H_2_O with 5% ACN and 0.125% FA) and auto-sampler loaded with an UltiMate 3000 HPLC pump onto a vented Acclaim Pepmap 100, 75 μm x 2 cm, nanoViper trap column (Thermo Fisher Scientific, Cat# 164535) coupled to a nanoViper analytical column (Thermo Fisher Scientific, Cat# 164570, 3 μm, 100 Å, C18, 0.075 mm, 500 mm) with stainless steel emitter tip assembled on the Nanospray Flex Ion Source with a spray voltage of 2000 V. An Orbitrap Fusion (Thermo Fisher Scientific) was used to acquire all the MS spectral data. Buffer B contained 99.875% ACN with 0.125% FA. The chromatographic run was for 4 h in total with the following profile: 0–7% for 7 min, 10% for 6 min, 25% for 160 min, 33% for 40 min, 50% for 7 min, 95% for 5 min and again 95% for 15 min respectively.

We used a multiNotch MS^3^-based TMT method to analyze all the TMT samples (McAlister et al., 2014; Ting et al., 2011; Weekes et al., 2014). The scan sequence began with a MS^1^ spectrum (Orbitrap analysis, resolution 120,000, 400–1400 Th, AGC target 2×10^5^, maximum injection time 200 ms). MS^2^ analysis, ‘Top speed’ (2 s), Collision-induced dissociation (CID, quadrupole ion trap analysis, AGC 4×10^3^, NCE 35, maximum injection time 150 ms). MS^3^ analysis, top ten precursors, fragmented by HCD prior to Orbitrap analysis (NCE 55, max AGC 5×10^4^, maximum injection time 250 ms, isolation specificity 0.5 Th, resolution 60,000).

MS^2^ CID-scan parameters were as follows; ion transfer tube temp = 300°C, Easy-IC internal mass calibration, default charge state = 2 and cycle time = 3 s. Detector type set to Orbitrap, with 60K resolution, with wide quad isolation, mass range = normal, scan range = 300–1500 m/z, max injection time = 50 ms, AGC target = 200,000, microscans = 1, S-lens RF level = 60, without source fragmentation, and datatype = positive and centroid. MIPS was set as on, included charge states = 2–6 (reject unassigned). Dynamic exclusion enabled with n = 1 for 30 s and 45 s exclusion duration at 10 ppm for high and low. Precursor selection decision = most intense, top 20, isolation window = 1.6, scan range = auto normal, first mass = 110, collision energy 30%, CID, Detector type = ion trap, OT resolution = 30K, IT scan rate = rapid, max injection time = 75 ms, AGC target = 10,000, Q = 0.25, inject ions for all available parallelizable time (Y.-Z. Wang et al., 2024).

### Mass spectrometry data processing

Raw spectra output files were converted using RawConverter (http://fields.scripps.edu/downloads.php), into MS1, MS2, and MS3 files. Files were uploaded to the Integrated Proteomics Pipeline (IP2; http://www.integratedproteomics.com) for protein identification and TMT quantification using ProLuCID, DTASelect, Census and Quantitative Analysis. Databases for MS searches included the protein sequences in the ordered library (i.e Metagenomic 1, Metagenomic 2, Protein Family), *E. coli* BL21(DE3) proteome downloaded from Uniprot, and dataset of common contaminants in proteomics (cRAP) downloaded from https://www.thegpm.org/crap/. The databases contained reverse sequences (decoys) analogous to each protein in the target database, used to estimate false discovery rate (FDR) and adjust for incorrect matches. ProLuCID search parameters include the amino acid modifications; 57.02146 C for carbamidomethylation and 304.2071 K for TMT 16-plex tagging, and peptide N-terminus differential modifications; 42.0106 for acetylation and 304.2017 for TMT 16-plex tagging. A fragment mass tolerance of 600 ppm and precursor delta mass cutoff of 5 ppm. Advanced search parameters utilized the DTASelect FDR type, requiring a minimum peptide length of six amino acid residues, minimum number of one peptide per protein, and minimum one tryptic end per peptide, with an estimated FDR of <1% for peptide matches. TMT-MS samples were quantified with IP2 by summing the tagged reporter ions.

### Calculating aggregation propensity

For each protein in each condition, we averaged the normalized TMT intensity across all 3 replicates. We then normalized post-stress intensities to the control sample by taking the ratio of the mean normalized TMT intensity in the post-stress condition to that of the control. Subsequently, we calculated aggregation propensity as the log_2_ of this fold change. Because the control samples and stress samples were not paired, the standard deviation was estimated using the propagation of uncertainty for ratios.

#### Correction for apparent increase in abundance

While proteins cannot truly increase in abundance after stress, some aggregation propensity measures appear positive due to TMT-MS normalization, in which equal amounts of peptides are analyzed for each condition. As a result, some proteins seem to increase to compensate for the loss of soluble protein caused by aggregation. To account for this effect, we normalized each dataset by centering the peak of the log_2_(fold change) distribution for each stress condition at zero. This adjustment was based on the observation that most proteins do not aggregate and facilitated comparison across conditions. In a small fraction of the proteins, even after this correction, measured increases exceeded the loss of total protein measured by BCA assay. We assumed that no single protein could increase more than the total protein loss so we clipped the upper limit to the maximum increase based on the BCA assay (Fig. S4).

#### Correction for library context effects

To assess how library context influenced measured aggregation, we included 1,050 identical proteins in both libraries, Metagenomic 1 and Metagenomic 2. Under acidic stress, we observed an offset between aggregation values for the overlapping proteins in Metagenomic 1 versus Metagenomic 2, with overall higher aggregation measured in Metagenomic 2. Therefore to correct for this systemic difference, we used the line of best fit determined based on the overlapping proteins (Eq.1) to adjust all aggregation measurements (log_2_(FC)) in the library Metagenomic 1, allowing data from both libraries to be combined for downstream analyses.

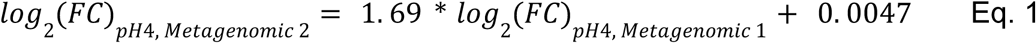

For the overlapping proteins, which had measurements in both libraries, we report the mean aggregation propensity — using the overlap-corrected Metagenomic 1 and uncorrected Metagenomic 2 measurements for pH stress, and uncorrected values for temperature stresses.

### Machine learning

#### Data filtering

Proteins that were not quantified in all control replicates were removed from the dataset. We aimed to investigate the properties that make globular proteins aggregation resistant so we removed any proteins that had a radius of gyration divided by the square root of protein length greater than 1.8 (Fig. S2), determined based on the lowest ranked AlphaFold 2-predicted structure.

#### Structure prediction

We used AlphaFold 2 (Jumper et al., 2021) to predict the protein structure for each protein in the libraries. We did not generate a multiple sequence alignment (MSA) for *de novo* designed sequences. We predicted 5 structures and then used the lowest ranked structure (highest pLDDT) for downstream analysis.

#### Feature extraction

We generated a set of interpretable protein features based on the protein sequence and structure. Structure-based features (e.g., radius of gyration or solvent exposed surface area) were calculated based on the lowest rank AlphaFold structure refined by Rosetta Relax protocol (Conway et al., 2014). Many protein features were generated using the pipeline from Ferrari et. al. 2025, Rosetta-based features from Rocklin et al. 2017, and prediction of intrinsic disorder from flDPnn (Hu et al., 2021). Additionally, we added more hydrophobicity and charge related features. We calculated sequence-based hydrophobicity using different scales from Expasy ProtScale (Aboderin, 1971; Abraham & Leo, 1987; Bull & Breese, 1974; Eisenberg et al., 1984; FAUCHERE & V, 1983; Guy, 1985; Hopp & Woods, 1981; Janin, 1979; Kyte & Doolittle, 1982; Manavalan & Ponnuswamy, 1978; Miyazawa & Jernigan, 1985; Mohana Rao & Argos, 1986; Monera et al., 1995; Parker et al., 1986; Roseman, 1988; Sweet & Eisenberg, 1983; Tanford, 1962; Welling et al., 1985; Wilson et al., 1981; Wolfenden et al., 1981). We also calculated a metric for hydrophobic clustering in the primary sequence, HpC, using the method from Bowman et al., 2020 with a sliding window size of 3 amino acids. We calculated isoelectric point and net charge based on the predicted structure using PROPKA3 (Olsson et al., 2011; Søndergaard et al., 2011) and based on the primary sequence using Biopython (pKs from Bjellqvist et al., 1993, 1994; Cock et al., 2009). For per-residue features, we calculated the mean, standard deviation, minimum, and maximum and used these features in our machine learning analysis.

#### Train, validation, and test splits

To test generalizability, we clustered the sequences in our machine learning data set using Mmseqs2 (Steinegger & Söding, 2017) with a minimum sequence identity threshold of 0.3. We then assigned clusters to train, test, and validation splits so that 70.5% of the proteins are in the train set, 9.2% are in the validation set, and 20% are in the test set (Fig. S9).

#### Model training

For machine learning analysis we combined the Metagenomic libraries, but did not merge them with the protein family library because there was an insufficient number of overlap sequences between the three libraries to assess and correct for library-specific effects (Fig. S2C). Additionally the protein family library exhibited a very small range of high temperature aggregation across domains, limiting its utility for machine learning.

We trained ridge regression and XGBoost models using Scikit-learn (Pedregosa et al., 2011) with four types of input features: amino acid composition, the full feature set (∼1,600 sequence- and structure-derived features), reduced feature sets for interpretability, and pretrained SaProt embeddings (Su et al., 2024). Reduced feature sets were generated from the training and validation sets by clustering correlated features (Fig. S9A). We computed a distance matrix from the absolute feature correlations and then performed hierarchical clustering using SciPy (Virtanen et al., 2020), applying different dendrogram thresholds to define feature clusters. For each stress condition and threshold, we selected the feature most correlated with aggregation for the training and validation sets. Additionally, we used pretrained SaProt language model (Su et al., 2024, 2025) to generate embeddings for each protein based on their structurally aware sequences using structure tokens generated from FoldSeek (van Kempen et al., 2024), based on their AlphaFold 2 predicted structures. Ridge regression and XGBoost models were trained with hyperparameters optimized on the combined training and validation data and evaluated on the held-out test set. For XGBoost models hyperparameters (e.g. max depth, learning rate, number of estimators) were optimized using randomized search with three-fold cross-validation and 20 iterations per model, minimizing mean squared error on validation folds. Ridge regression alpha parameters were optimized in the range of 10^-3^ to 10^3^. Model performance was reported as the mean ± half 95% confidence interval of the bootstrapped (n = 1000 samples) test set predictions. Separate models were trained to predict aggregation induced by 50 °C, 75 °C, and pH 4.

We fine-tuned the 650M-parameter SaProt language model (Su et al., 2024) with LoRA via SaProtHub (Su et al., 2025). Models were trained with structurally aware sequences as defined by FoldSeek (van Kempen et al., 2024) from the AlphaFold 2-predicted structure. Hyperparameters were optimized based on performance on the validation set; the final models used a learning rate of 5 × 10^-5^, LoRA alpha of 16, LoRA dropout of 0.1, and LoRA rank of 8. We saved the best checkpoint based on performance on the validation set and final model performance was evaluated on the held-out test set. We trained separate models to predict 50 °C, 75 °C, and pH 4-induced aggregation. All model performance metrics are represented as mean ± half 95% CI determined by bootstrapping the test set predictions (n= 1000 samples).

### Predictions with published tools

Aggregation and solubility predictions were performed using several sequence- and structure-based tools. TANGO (Fernandez-Escamilla et al., 2004) and CamSol (Sormanni et al., 2015) predictions were computed at both pH 7 and pH 4. ANuPP (Prabakaran et al., 2021) and PASTA (Trovato et al., 2007; Walsh et al., 2014) predictions were obtained using the default settings on their respective web servers. AgMata (Orlando et al., 2020) predictions were made with the Bio2Byte tools Python package. Aggrescan 3D (Conchillo-Solé et al., 2007) predictions were on the AlphaFold 2 predicted structures using the A3D python package (Kuriata et al., 2019). Spatial aggregation propensity (Chennamsetty et al., 2009) was in PyRosetta (Chaudhury et al., 2010) using the AlphaFold 2 predicted structure. Solubility predictions were obtained from the NetSolP-D model (distilled ESM1b model, Thumuluri et al., 2022). Predictions were also made with the 650M SaProt model fine-tuned on the DeepSol dataset (Khurana et al., 2018; Su et al., 2025) and on the Meltome thermostability dataset (Jarzab et al., 2020; Su et al., 2025). For tools providing per-residue scores (e.g., AgMata, Aggrescan 3D, and PASTA), we compared our experimental data to the mean predicted score across residues.

### Predictions on published test sets

We tested our 75 °C model predictions on published datasets measuring soluble protein expression (Niwa et al., 2009; Uemura et al., 2018), mutation effects on solubility (Attanasio et al., 2025), apparent melting temperature (Jarzab et al., 2020), amyloid properties of peptides (Louros et al., 2020), and amyloid nucleation properties of peptides (M. Thompson et al., 2025). For peptides, predictions were made using only the amino acid sequence, whereas for proteins we used the AlphaFold 2-predicted structure or experimentally determined PDB structures. Predictions for Uemura et al. (2018) were made on structurally aware sequences derived from AlphaFold predicted structures from AlphaFoldDB (Fleming et al., 2025; Jumper et al., 2021). Predictions for Niwa et al. 2009 used published structurally aware sequences from AlphaFold 2 predicted structures (Tan et al., 2025). Predictions for Attanasio et al. 2025 were made on structurally aware sequences based on published PDB or AlphaFold 2 predicted structures of the wild-type proteins, with mutant sequences generated by substituting the corresponding amino acids in the wild-type structural sequence. Two proteins from the SOulMuSiC dataset were excluded because their PDB codes did not match the provided structure file names. Predictions for the human-cell FLIP split (Dallago et al., 2022) from Jarzab et al. (2020) were made on published structurally aware sequences from the AlphaFold 2 predicted structure (Su et al., 2025). For WaltzDB peptides, entries annotated as both amyloid and non-amyloid were excluded from this analysis.

### Single substitution effects analysis

This analysis focused on proteins in the test set, specifically clusters containing at least five proteins. For each cluster, we generated multiple sequence alignments (MSA) using the MUSCLE algorithm (Edgar, 2004) in SeaView (Gouy et al., 2010). At each aligned position, we considered amino acid groups represented by at least three proteins, leaving a total of 61 clusters and 1,480 aligned positions in these clusters.

We then used our 75 °C SaProt model to predict deep mutational scans for all domains within the selected clusters. For each single mutant, we used the FoldSeek sequence from the wild-type structure and substituted the corresponding residue as inputs to the model. We did not generate AlphaFold 2 models for each mutant because (i) generating that many structures would be computationally expensive, (ii) it does not reliably predict structural changes for single substitutions, and (iii) for a small sample of the test set we saw minimal differences between using the predicted structure versus the wild-type structure (Fig. S15D). To reduce noise and account for amino acids with similar effects, we clustered the average predicted effect for each amino acid within each cluster and then grouped amino acids by their predicted impact on aggregation (Fig. S15C). The resulting amino acid groups were: P; DE; QNG; HKSAT; FW; ILY; R; MV (Fig. S15C).

Within protein clusters, we estimated the experimental effects of single point substitutions. For each aligned position, we grouped proteins with related amino acid identities according to the amino acid groups defined above. We then averaged the log_2_(fold change) for all proteins with a specific amino acid group at a specific position. We then calculated the difference between average log_2_(fold change) between each amino acid group pairs across the 1,480 positions in the 61 clusters.

Finally, within protein clusters, we then calculated the predicted substitutions effects between amino acid groups at each aligned position. For example, in cluster 105, we quantified the predicted effect of substituting amino acids I, L, or Y with either D or E at position 27 in the MSA. For the three wild-type proteins containing L at this position, we averaged the predicted Δlog_2_(fold change) for all six predictions (three wild-type proteins x two amino acids). We repeated this procedure for all amino acid group substitutions across the 1,480 positions in the 61 clusters.

## Data Availability

All data used to generate the results in this study are available to download at https://forms.gle/iJZFFXpiy81S3nfj7.

## Acknowledgements

We thank Kotaro Tsuboyama for the cDNA display stability proteolysis measurements used in this work. We thank Xiaoyu Zhang for insightful discussions, feedback, and assistance with mass spectrometry experiments. We thank Michal Zielinski and Rosalia Schneider for their assistance assembling the MGnify domain library. We thank members of the Rocklin Lab for their valuable discussions and comments on this manuscript. We thank the Northwestern Proteomics Core for their assistance with mass spectrometry experiments. This research was supported in part through the computational resources and staff contributions provided for the Quest high performance computing facility at Northwestern University which is jointly supported by the Office of the Provost, the Office for Research, and Northwestern University Information Technology. This work was supported by the National Institutes of Health (NIH) awards DP2GM140927 (G.J.R.), 1R35GM158118-01 (G.J.R.), and S10 OD032464 (J.N.S), the Northwestern Chemistry of Life Processes Training Grant NIH T32GM105538 (C.M.M.), the PhRMA Foundation predoctoral fellowship in drug delivery (C.M.M.), the Northwestern Chemistry of Life Processes CAURS Undergraduate Research Award (Y.M.G).

## Contributions

C.M.M. and G.J.R. designed the research. C.M.M. expressed and purified protein libraries and individual proteins and performed the aggregation experiments, MS sample preparation, data analysis, and machine learning. K.K.G. and H.M.V. performed mass spectrometry and data processing under the supervision of J.N.S. Y.M.G. cloned DNA constructs and M.D.J. purified individual proteins. G.J.R. acquired funding and supervised the project. C.M.M. and G.J.R. wrote and revised the manuscript with input from all of the authors.

## Supplemental Figures

**Figure S1.**
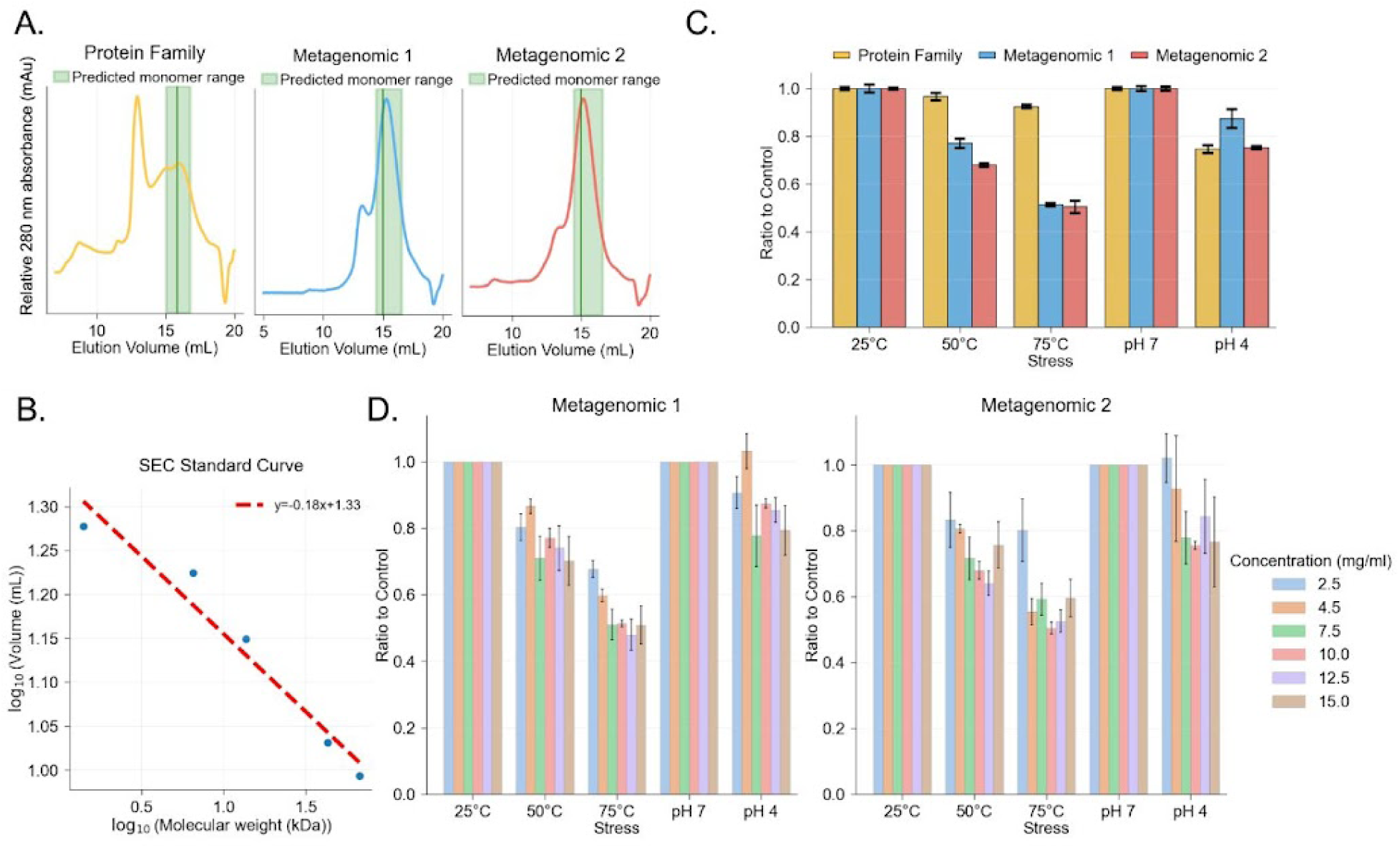
Aggregation assay samples and conditions. (A) Size exclusion chromatography traces for each purified library. The green area indicates the predicted monomeric fraction, determined using a standard curve of reference proteins. (B) Standard curve based on the SEC trace collected on our instrument for a standard set of proteins (BSA: 67 kDa, Ovalbumin: 43 kDa, Ribonuclease A: 13.7 kDa, Aprotinin: 6.5 kDa, Vitamin B12: 1.4 kDa). The linear fit was used to calculate expected monomer elution. (C) Total protein concentration in the supernatant after centrifugation, shown as a ratio relative to the control sample and measured by BCA assay for each stress condition. Libraries are colored as follows: Protein Family (yellow), Metagenomic 1 (blue), and Metagenomic 2 (red). (D) Total protein concentration in the supernatant after centrifugation across a range of starting concentrations for each stress condition, shown as a ratio relative to the control sample and measured by BCA assay. Data are shown for Metagenomic 1 (left) and Metagenomic 2 (right).

**Figure S2.**
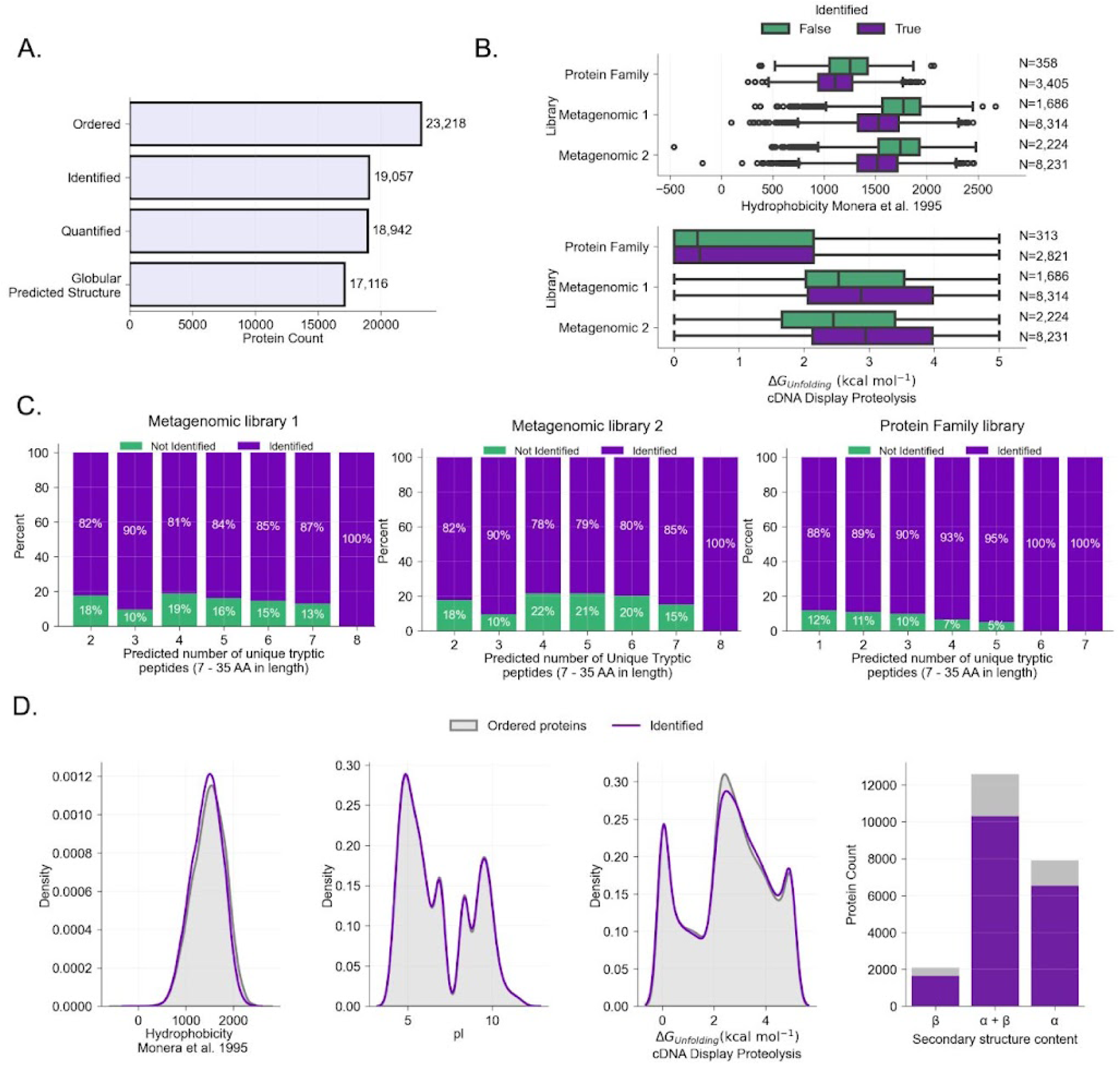
Comparison of proteins identified and not identified by MS. (A) Counts of unique protein domains that were ordered, identified by mass spectrometry, quantified, and determined to have a globular predicted structure. Counts combine the Metagenomic 1, Metagenomic 2, and Protein Family libraries and represent the number of unique domains. (B) Distribution of hydrophobicity and ΔG_unfolding_ measured by cDNA display proteolysis for identified versus not identified proteins in each library. Fewer proteins are shown in the Protein Family library for ΔG_unfolding_ because some proteins were not assayed with cDNA display proteolysis. (C) Percentage of proteins identified (purple) or not identified (green) based on the predicted number of unique tryptic peptides. Data are shown separately for each library: Metagenomic 1 (left), Metagenomic 2 (middle), Protein Family (right). (D) Protein feature distributions for all ordered protein domains versus the identified proteins. Hydrophobicity was calculated using the scale from Monera et al. (1995), based on reversed-phase HPLC retention times of amphipathic α-helical peptides with single amino acid substitutions.

**Figure S3.**
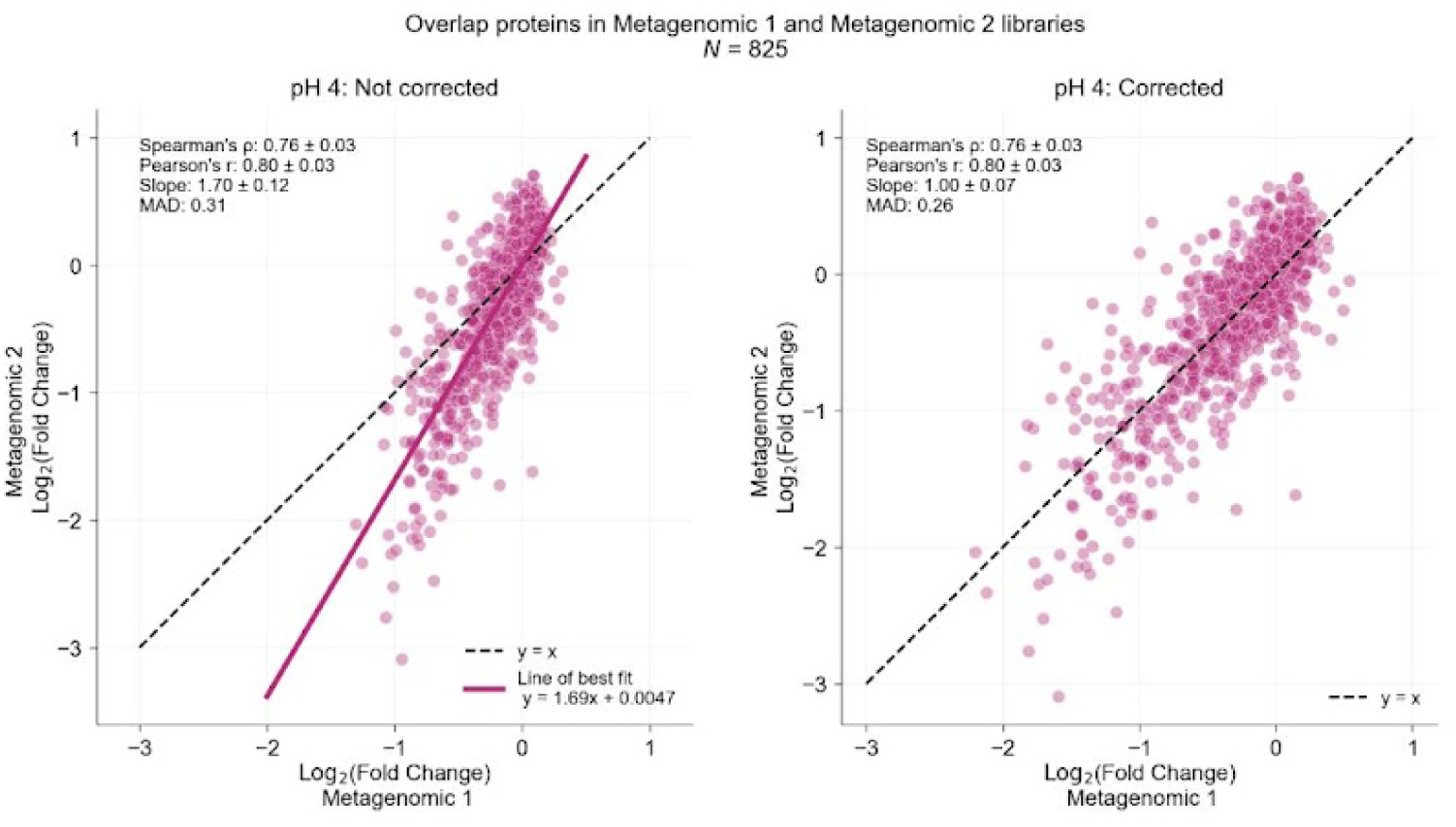
Correction of acid-induced aggregation using overlap proteins. Correlation of acid-induced aggregation propensity measured for 825 proteins in Metagenomic 1 and Metagenomic 2 libraries (left) and comparison of the corrected Metagenomic 1 values to the Metagenomic 2 measurements (right). The black dotted line indicates y=x and the solid pink line indicates the line of best fit. The Metagenomic 1 values were corrected based on the line of best fit. Metrics are represented as values ± half 95% CI from bootstrapping.

**Figure S4.**
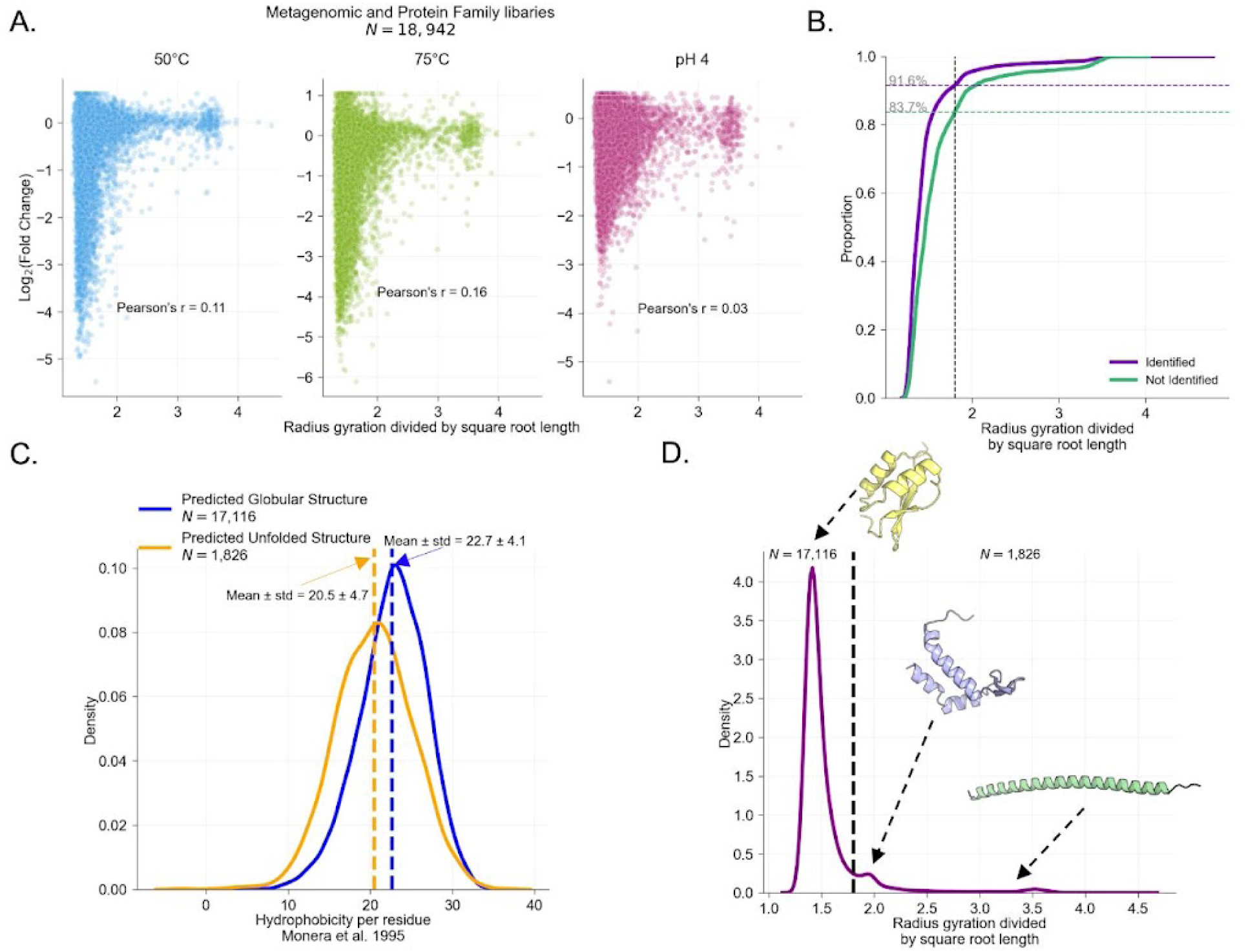
Filtering proteins with globular predicted structures. (A) Correlation between radius of gyration and aggregation in different stress conditions. The radius of gyration was calculated using the lowest rank AlphaFold-predicted structure and normalized by the square root of sequence length (number of residues) to remove length dependence. (B) Comparison of the radius of gyration for identified (purple) and not identified (green) proteins for all domains in the Metagenomic and Protein Family libraries. The gray line shows a cutoff of 1.8, where proteins below this cutoff show a more globular structure. The colored dotted lines and grey label indicate the proportion of proteins in each category that fall below the cutoff. (C) Hydrophobicity distributions for proteins with a predicted globular structure (blue, radius of gyration divided by square root of length <= 1.8) and a predicted unfolded structure (orange, radius of gyration divided by square root of length > 1.8). The dotted lines show the mean ± standard deviation for each distribution. Hydrophobicity was calculated using the scale from Monera et al. (1995), based on reversed-phase HPLC retention times of amphipathic α-helical peptides with single amino acid substitutions. (D) Distribution of radius of gyration divided by the square root of protein length for all quantified domains. We set the cutoff at 1.8 to select only globular proteins for our machine learning analysis. Representative AlphaFold structures for proteins with different radii of gyration are shown.

**Figure S5.**
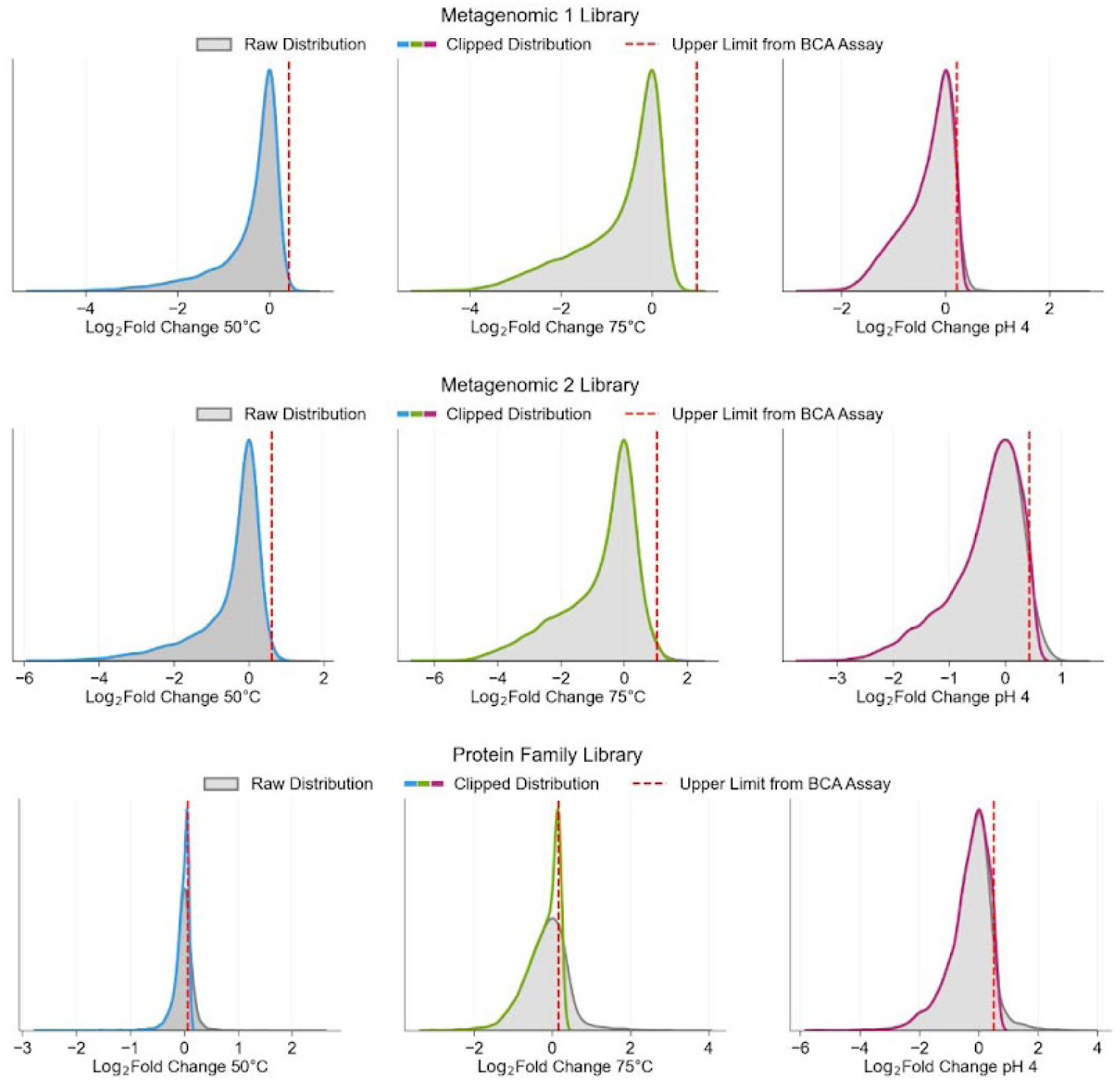
Clipping aggregation metric based on total protein loss. Each panel shows the distribution of log_2_(fold change) (grey distribution, peak centered at 0) and the maximum upper limit a single protein could increase based on the total protein loss calculated by the BCA assay (red line) for each stress condition. The distribution was clipped at the upper limit (colored line) and these are the aggregation measures analyzed throughout the paper. Each panel shows the correction for a different library (top: Metagenomic library 1, middle: Metagenomic library 2, bottom: Protein Family library).

**Figure S6.**
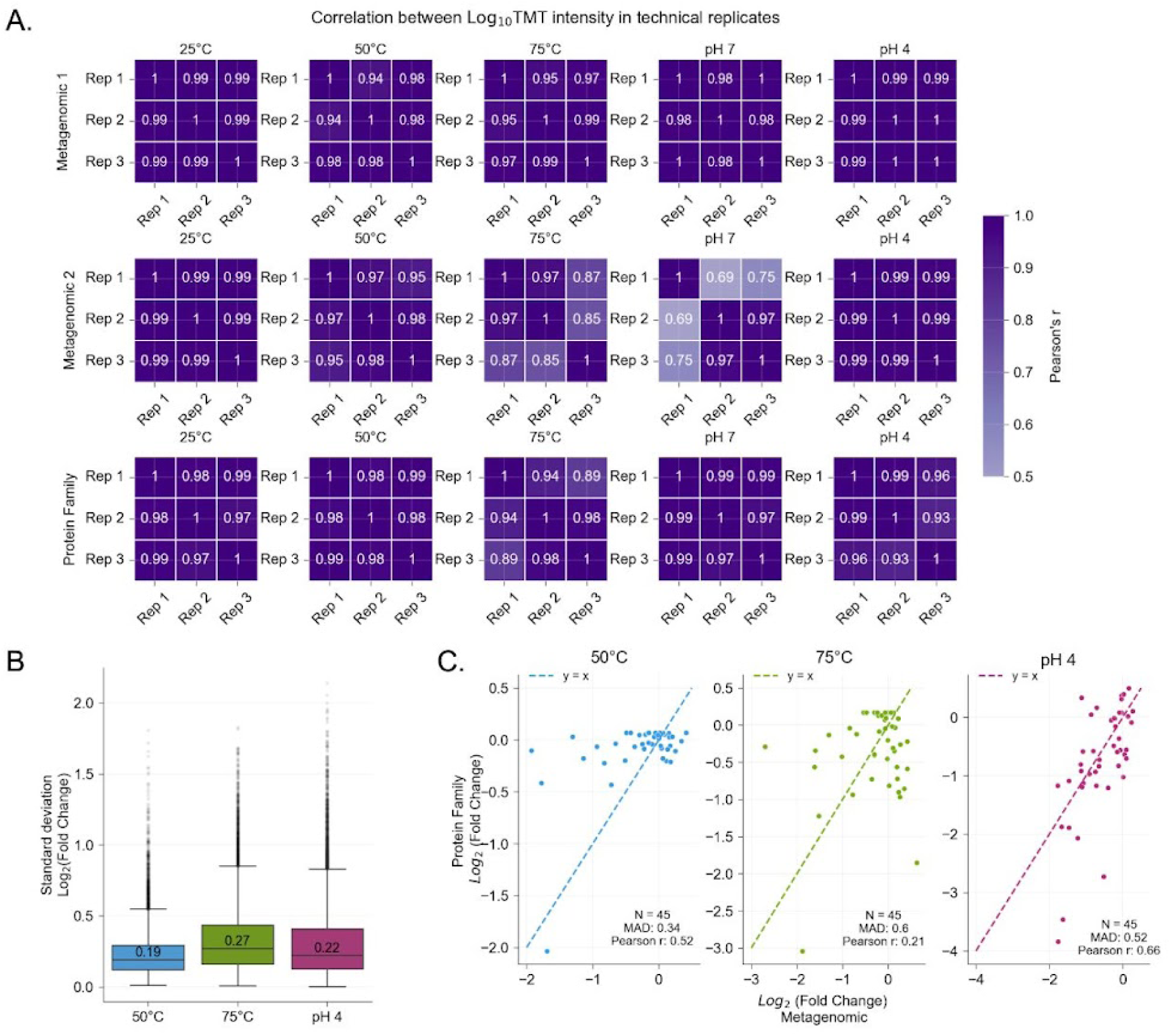
Reproducibility of aggregation measures. (A) Pearson’s correlation matrix between normalized MS intensity in replicate samples for each stress condition in each library. (B) Distribution of standard deviation in log_2_(fold change) in each aggregation stress. The median is written on the boxes. Boxplots show the median (center line), 25th and 75th percentiles (box edges), and whiskers extending to 1.5x the interquartile range. (C) Comparison of aggregation measured in the metagenomic library versus protein family library for 45 proteins quantified in both libraries (out of a set of 50 proteins assayed in both libraries), using the corrected pH 4 log_2_(fold change) values for the Metagenomic libraries.

**Figure S7.**
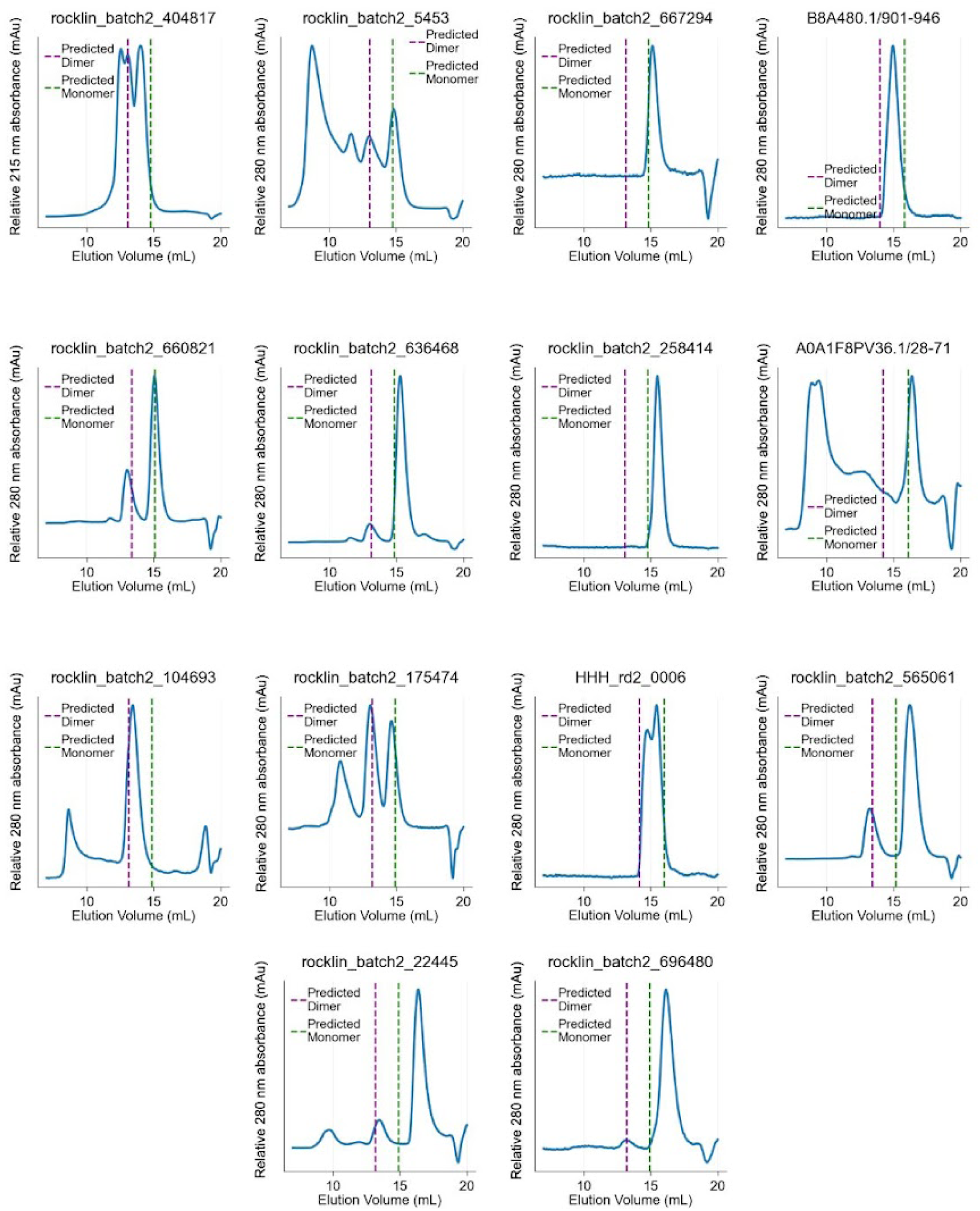
Size exclusion chromatography of purified individual proteins. Chromatograms show absorbance at 280 nm for each protein, except for rocklin_batch2_404817, which shows the absorbance at 215 nm because this protein lacks tryptophan and tyrosine residues. The purple and green lines indicate the expected elution volumes for dimers and monomers, respectively, based on a standard curve (Fig. S1B).

**Figure S8.**
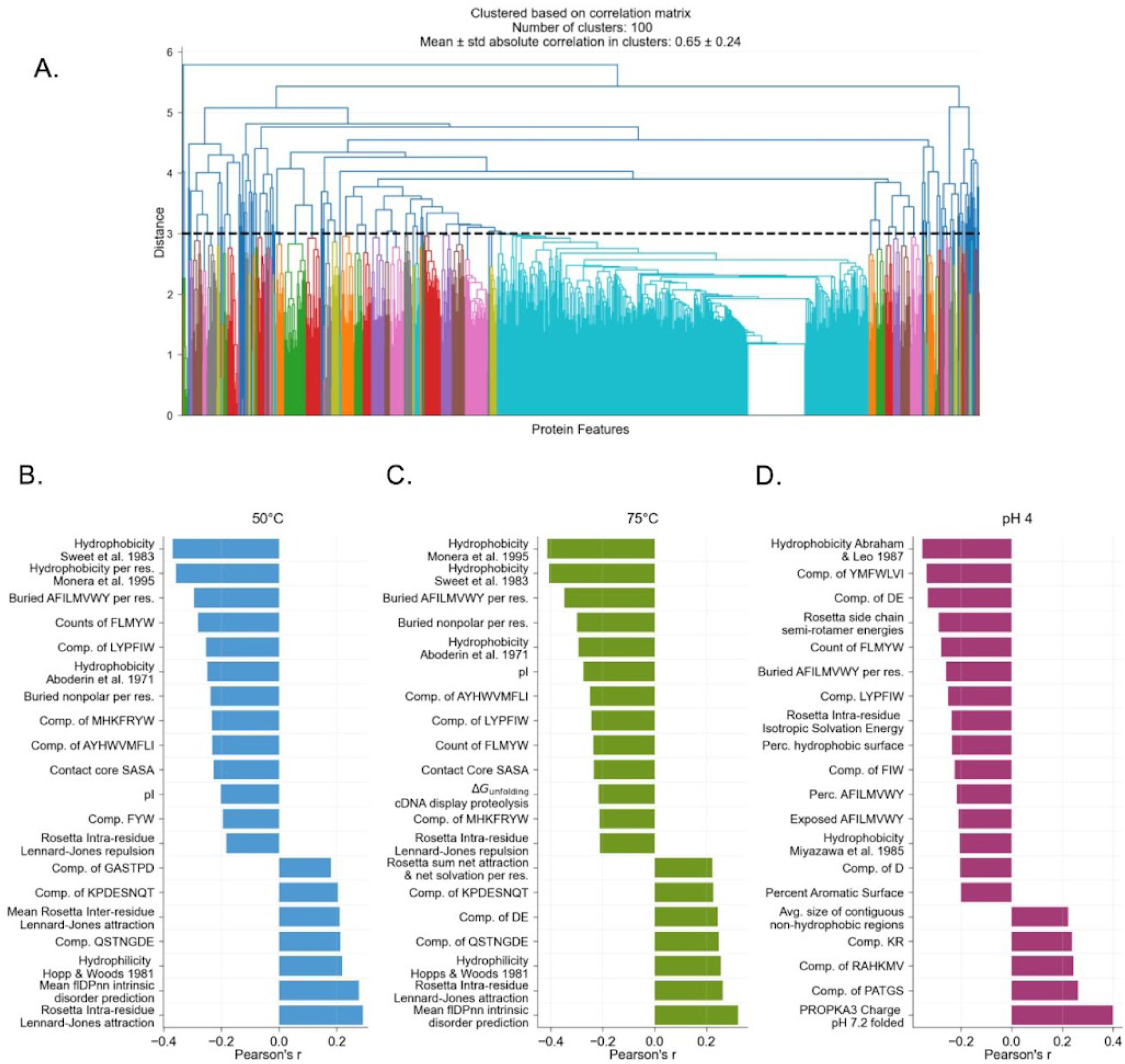
Top correlated protein features with aggregation. (A) Hierarchical clustering of protein features based on the correlation matrix. Clusters were defined using a dendrogram threshold of 3, which grouped the features into 100 distinct clusters. (B–D) The most strongly correlated feature from the top 20 feature clusters associated with aggregation under each condition: (B) 50 °C (blue), (C) 75 °C (green), and (D) pH 4 (pink). Displayed hydrophobicity scales include: Sweet et al. 1983, based on the free energy cost of moving each amino acid from a hydrophobic to hydrophilic environment; Monera et al., 1995, based on reversed-phase HPLC retention times of amphipathic α-helical peptides with single amino acid substitutions; Hopp & Woods 1981, hydrophilicity scale derived from a hydrophobicity scale (Nozaki & Tanford, 1971); Abraham & Leo 1987, based on fragment-based calculations of amino acid side-chain partition coefficients; Miyazawa et al. 1985, based on the contact energies derived from observed residue interactions in 3D protein structures.

**Figure S9.**
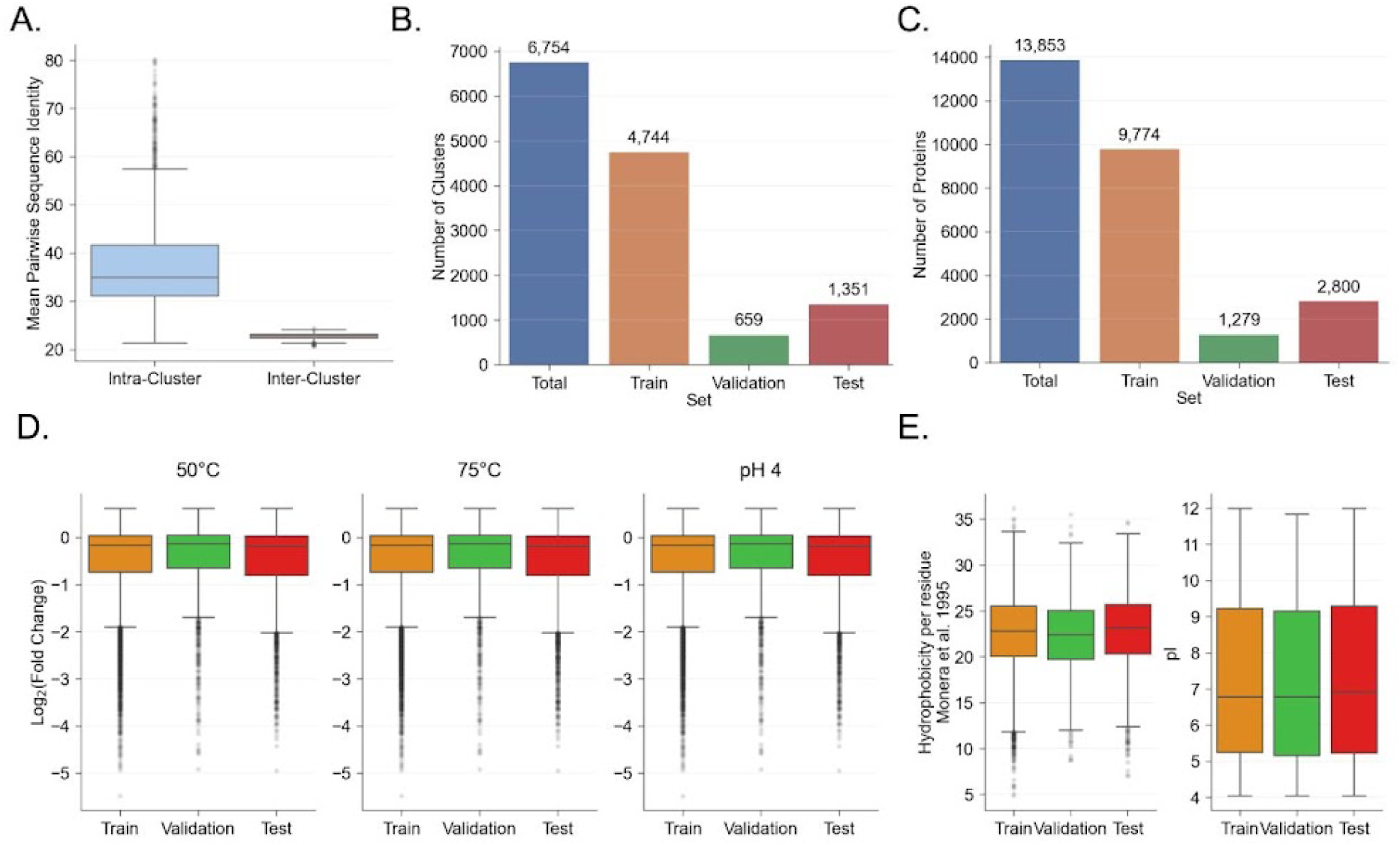
Clustering and splitting into train, test, and validation sets. (A) Mean pairwise sequence identity for pairs in the cluster (Intra-Cluster, blue) and pairs between proteins in the cluster and not in the cluster (Inter-Cluster, orange). (B) Number of clusters in the dataset and each split. (C) Number of proteins in the dataset and each split. (D) Distribution of each aggregation phenotype in the train, validation, and test set. (E) Distribution of hydrophobicity and pI for the proteins in the train, validation, and test set. Hydrophobicity scale from Monera et al., 1995, based on reversed-phase HPLC retention times of amphipathic α-helical peptides with single amino acid substitutions. pI was calculated using pK values from Bjellqvist et al. (1993 & 1994). The boxplots show the median (center line), 25th and 75th percentiles (box edges), and whiskers extending to 1.5x the interquartile range.

**Figure S10.**
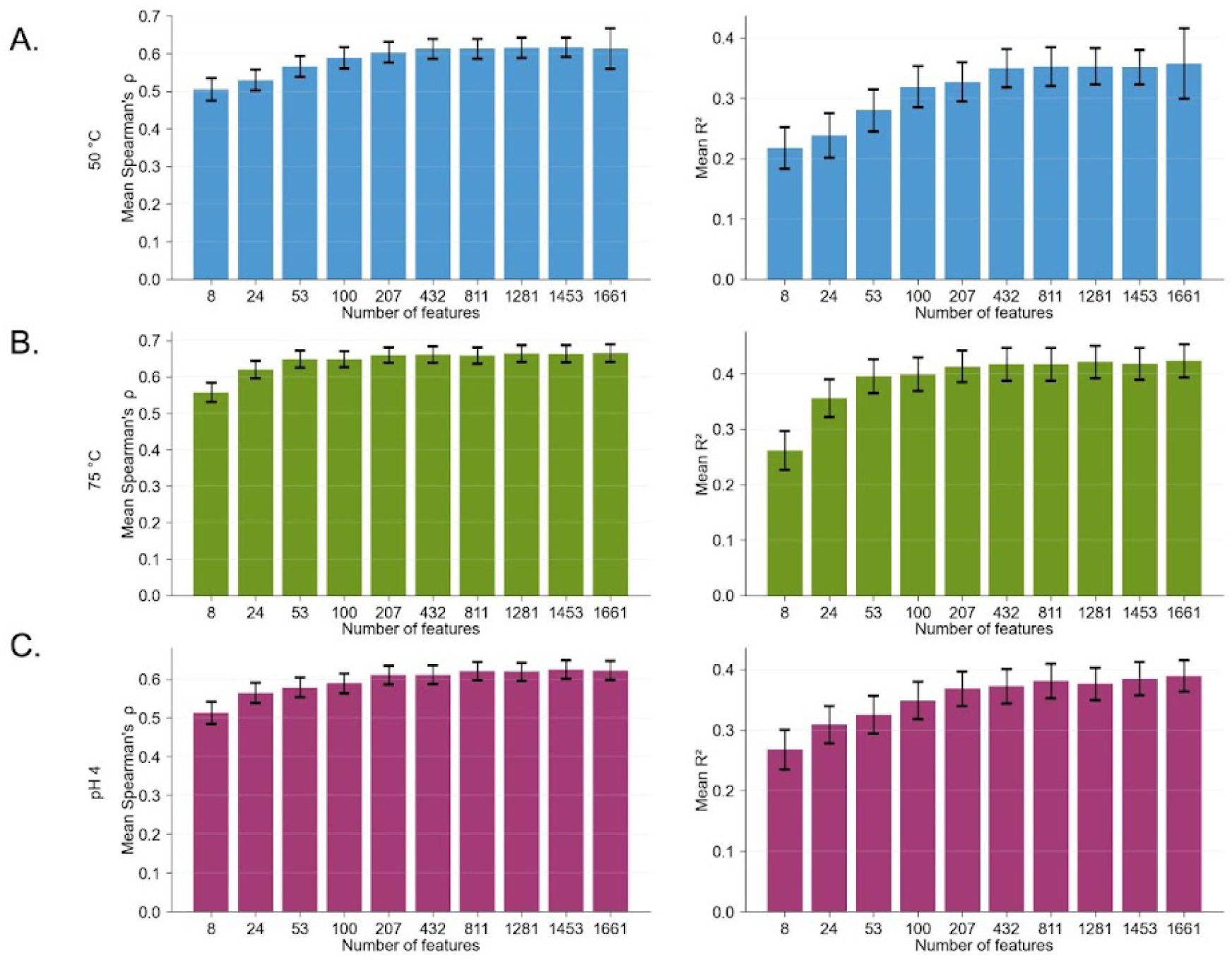
Performance of XGBoost models with different numbers of input features. Each panel shows model performance for the different aggregation stresses: (A) 50 °C, (B) 75 °C, (C) pH 4. The models were trained on different sets of features determined using different thresholds for clustering protein features on their correlation matrix (Fig. S8A). Once features were clustered, we selected the feature with the highest correlation to the target variable in the combined training and validation set for each cluster. The left plots show the mean Spearman’s correlation and the right shows the mean R^2^ from bootstrapping. Error bars represent the half 95% CI from bootstrapping.

**Figure S11.**
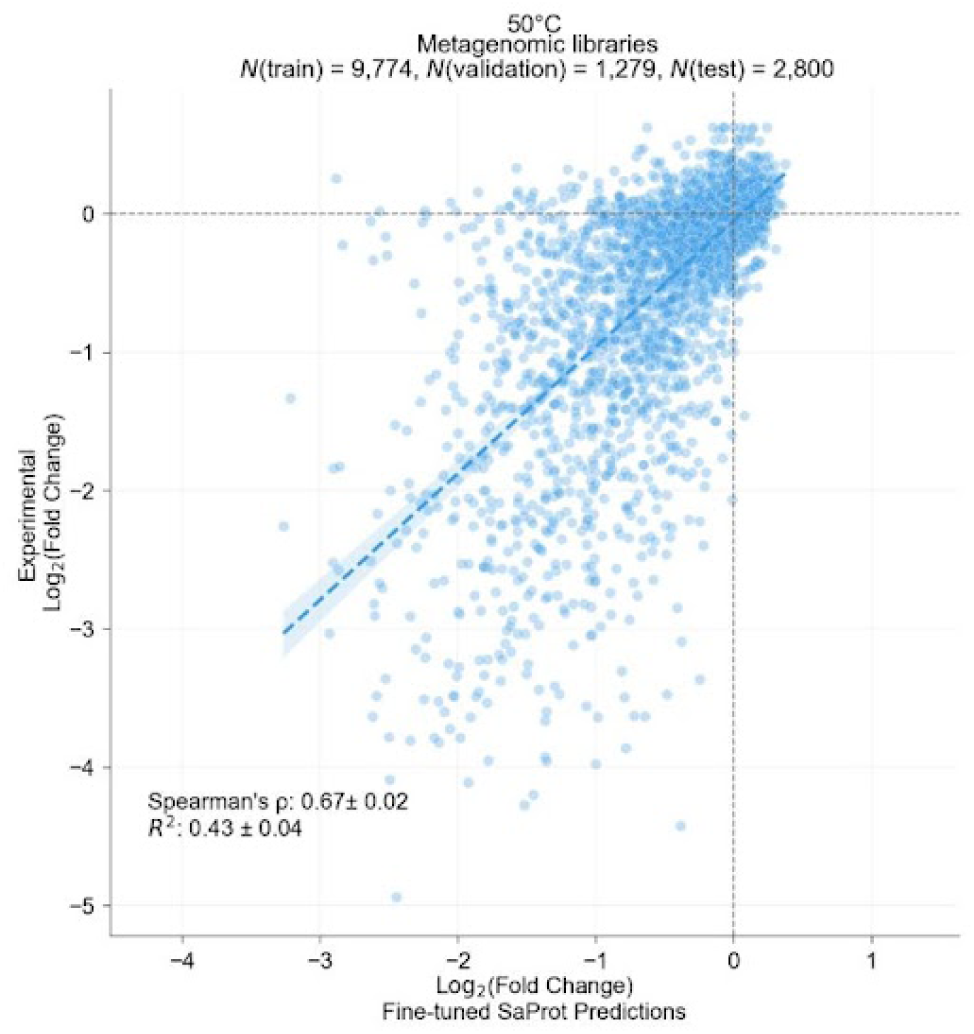
Finetuned SaProt predictions versus experimental values for 50 °C aggregation on the held-out test set. Statistics are reported as mean ± half 95% CI from bootstrapping.

**Figure S12.**
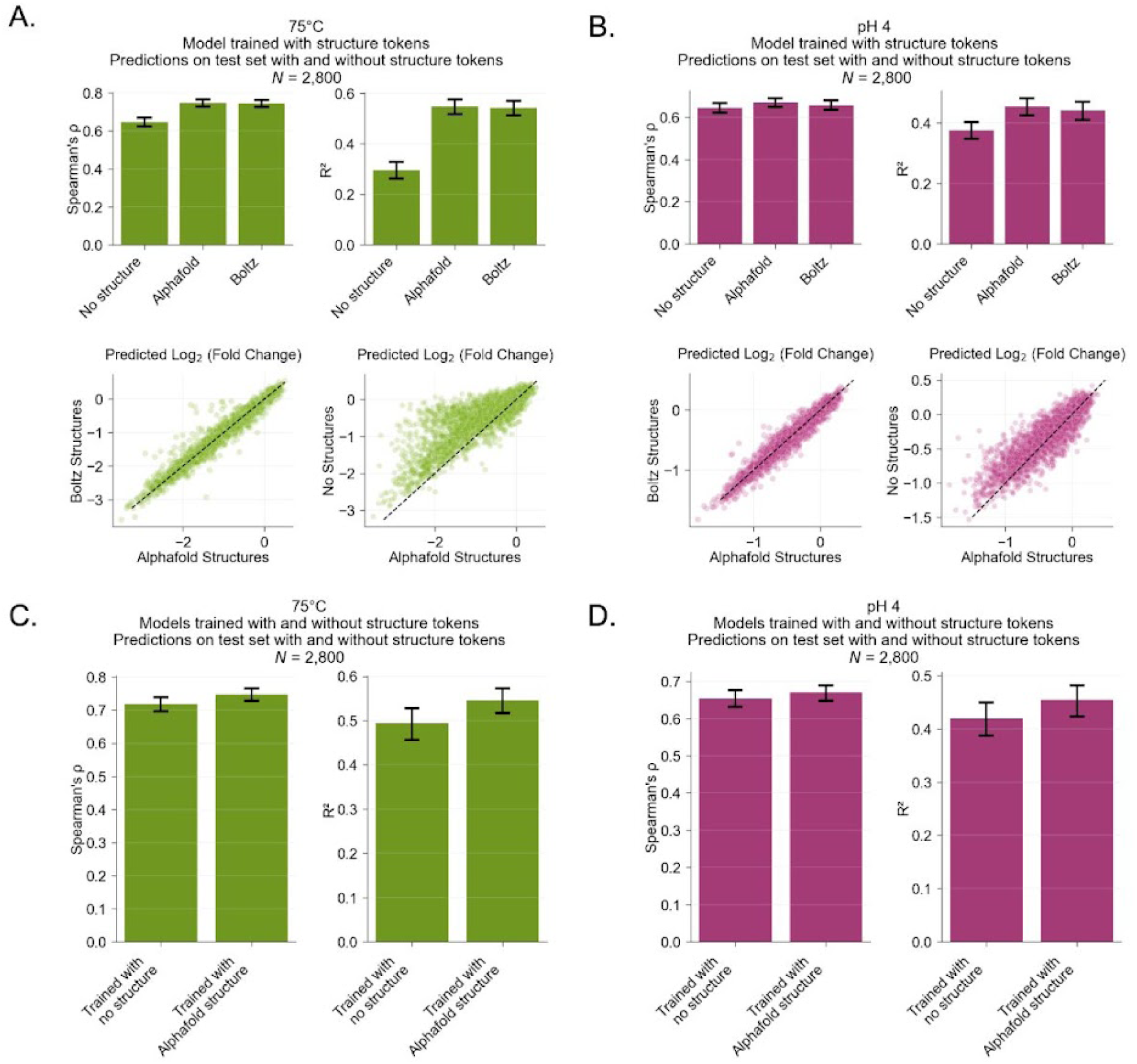
SaProt predictions with and without structural information. (A) Model performance for predicting aggregation induced by 75 °C stress using different structural token inputs. (B) Model performance for acid-induced aggregation using different structural token inputs. Structures were predicted using either AlphaFold 2 (Jumper et al., 2021) or Boltz (Passaro et al., 2025), and Foldseek (van Kempen et al., 2024) was used to generate structurally aware sequences for SaProt predictions. For predictions without structural information, we used amino acid sequences with all structural tokens masked. In panels A-B, the same fine-tuned SaProt models (trained on AlphaFold 2 predicted structures) were used to predict aggregation across the three test set versions. Panels D-E compare SaProt models either trained on structurally aware sequences with structure tokens from AlphaFold 2 predicted structures or trained only on amino acid sequences without structure tokens. (D) Performance for predicting aggregation induced by 75 °C stress with these two models. (E) Performance for acid-induced aggregation with these two models. In all bar plots, bars represent the mean and error bars represent half of the 95% CI estimated by bootstrapping.

**Figure S13.**
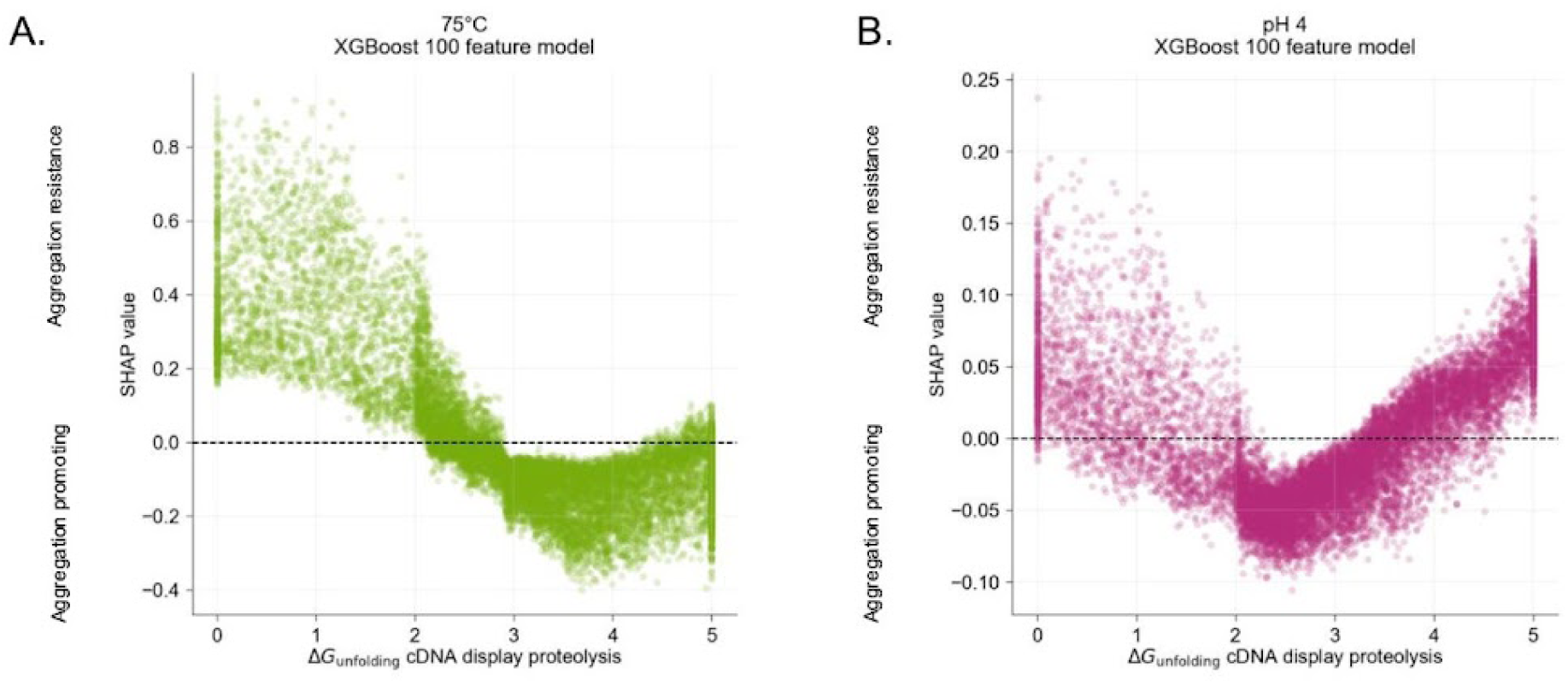
SHAP values for ΔG_unfolding_ in predicting aggregation using XGBoost models. (A) SHAP values for ΔG_unfolding_ from models trained to predict 75 °C-induced aggregation plotted against ΔG_unfolding_. (B) SHAP values for ΔG_unfolding_ from models trained to predict acid-induced aggregation versus ΔG_unfolding_. Higher SHAP values indicate that ΔG_unfolding_ contributes to aggregation resistance, whereas lower values indicate that ΔG_unfolding_ is aggregation promoting. Each point represents the feature value for a protein in the training or test set. SHAP values are calculated from the XGBoost models trained on 100 protein features.

**Figure S14.**
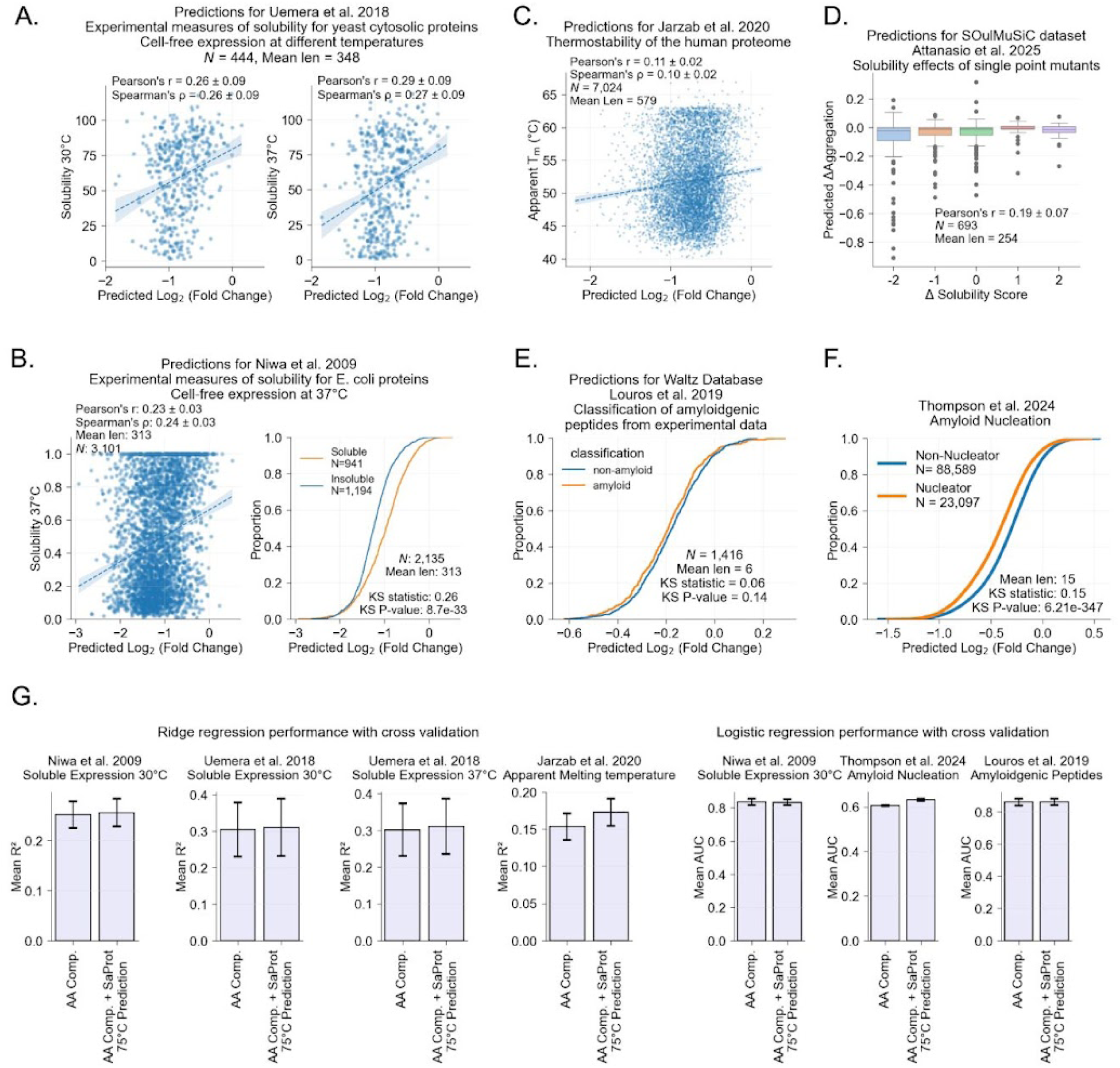
SaProt 75 °C predictions on published datasets of similar phenotypes. (A) Predicted 75 °C-induced aggregation versus experimental solubility of yeast cytosolic proteins expressed at 30 °C (left) or 37 °C (right) (Uemura et al., 2018). (B) Predicted 75 °C-induced aggregation versus soluble expression of *E. coli* proteome at 37 °C (Niwa et al., 2009). (C) Predicted 75 °C-induced aggregation versus experimentally measured thermostability (apparent melting temperature, T_m_) across the human proteome (Jarzab et al., 2020). Thermostability data from the FLIP task using structurally aware sequences (Su et al., 2025). (D) Predicted changes in aggregation (ΔAggregation at 75 °C) versus experimentally measured solubility changes (ΔSolubility Score) for mutants and associated wild types in the SOuLMuSiC dataset (Attanasio et al., 2025). (E) Predicted 75 °C-induced aggregation for peptides experimentally classified as amyloidogenic (orange) or non-amyloidogenic (blue) (Louros et al., 2020). (F) Predicted 75 °C-induced aggregation for peptides experimentally classified as amyloid nucleators (orange) or non-nucleators (blue) (M. Thompson et al., 2025). (G) Ridge regression (left) and logistic regression (right) models trained with cross-validation to predict published datasets. Models were trained with either amino acid composition only or amino acid composition plus 75 °C SaProt predictions. Bars and error bars represent the mean ± half 95% CI from bootstrapping the predictions across folds. In panels A–C, Pearson’s *r* and Spearman’s ρ are reported as mean ± half 95% CI from bootstrapping. In panels E–F, distributions were compared using the two-sample Kolmogorov–Smirnov (KS) test.

**Figure S15.**
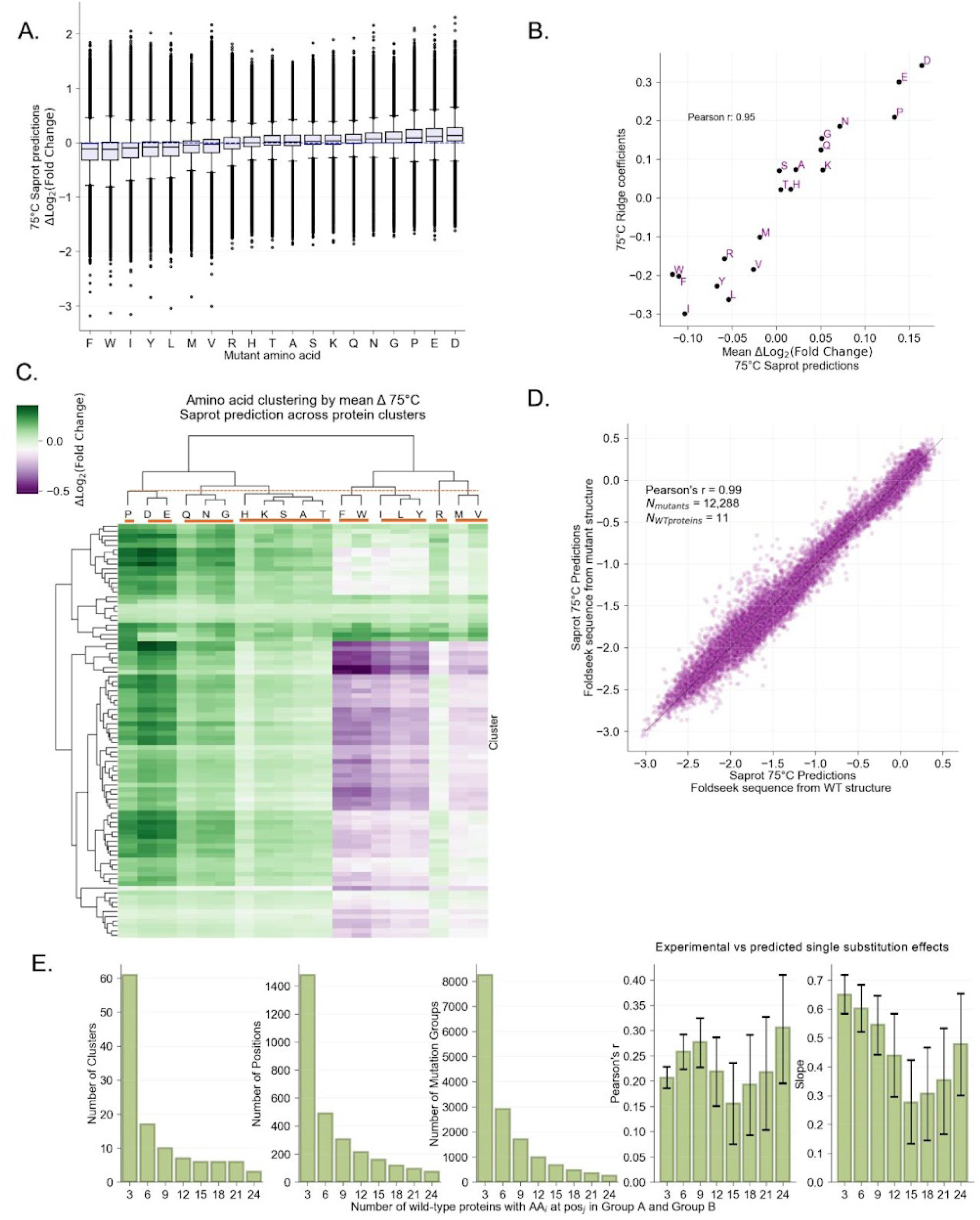
Distribution, clustering, and validation of SaProt-predicted amino acid substitution effects for 75 °C aggregation. (A) Distribution of Δ SaProt substitution predictions for each amino acid. ΔLog_2_(Fold Change) is defined as mutant prediction minus wild-type prediction; more negative values indicate that mutating to that residue is predictive of increased aggregation propensity. The blue dotted line indicates 0. The boxplots show the median (center line), 25th and 75th percentiles (box edges), and whiskers extending to 1.5x the interquartile range. (B) Comparison of SaProt predicted amino acid substitution effects and ridge regression coefficients from the amino acid composition model. (C) Clustering of amino acids based on the predicted effects of mutations to each residue. Each row indicates a cluster, colored by the mean Δlog_2_(Fold Change). The orange dotted line indicates the clustering threshold and the solid orange lines delineate amino acid groups. (D) Comparison of SaProt 75 °C predictions for a set of mutant proteins using structure token from either the wild-type predicted structure or the mutant predicted structure. (E) Comparison of experimental differences between wild types containing group A vs group B amino acids at a given position, and fine-tuned SaProt predicted effects for mutations from group A to group B for different cutoffs of the number of proteins in group A and group B. Analyses were performed within test set clusters with ≥ 5 proteins. Pearson’s r and slope were calculated by comparing the predicted mutation effect (group A→ group B) to the experimental effect (AA in group A vs group B). Slope and Pearson’s r values indicate mean ± half 95% CI from bootstrapping.

**Table S3.**
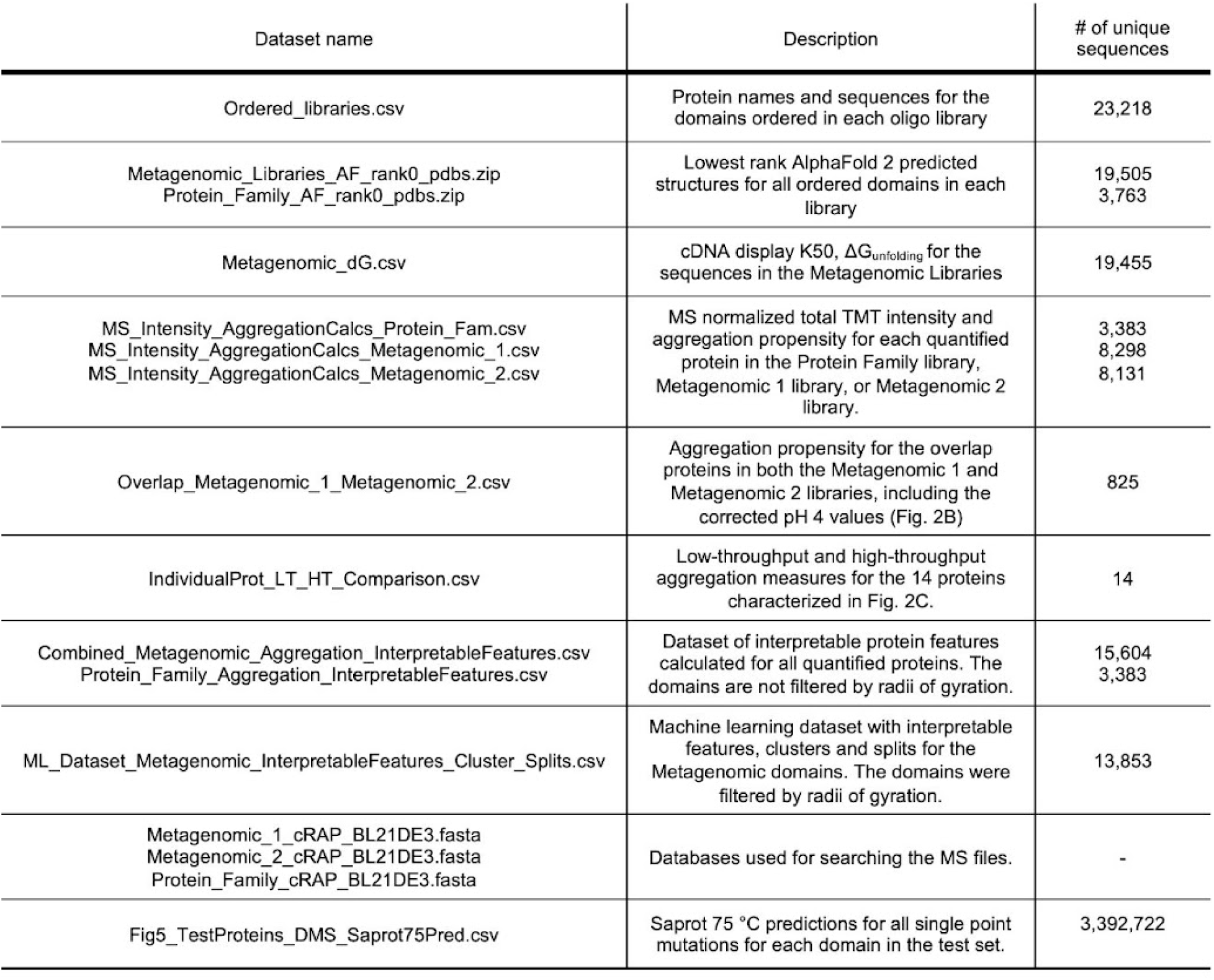
Published datasets available at: https://forms.gle/iJZFFXpiy81S3nfj7.

